# EFFECTS OF SUBCHRONIC EXPOSURE TO POLYSTYRENE NANOPLASTICS ON THE MOUSE SMALL INTESTINE

**DOI:** 10.64898/2026.09.15.751754

**Authors:** Mary Gregory, Julie Dufresne, Paloma Da Cunha De Medeiros, Unnikrishnan Kannan, Sara Yim, Pauline Moinard, François Gagné, Stéphane Lefrancois, Daniel G. Cyr

**Author notes:** Address for Correspondence: Dr. Daniel Cyr, INRS-Centre Armand-Frappier Santé Biotechnologie, Université du Québec, 531 boulevard des Prairies, Laval, QC, Canada, H7V1B7, (ext 8833).

## Abstract

Nanoplastics (NPs) are being used increasingly in cosmetics, personal care products, foods, automotive products, and cleaning products, as well as being by-products of some industrial processes. The aim of this study was to examine the effects of a subchronic exposure of NPs on various biological systems, using mice as a model. Adult male mice were administered 500 nm polystyrene (PS) NPs at 0.15 mg/day and 1.5 mg/day for 60 days, to mimic subchronic exposure to NPs. Control mice received water. The small intestines were separated into three regions: duodenum, jejunum, and ileum. Hematoxylin and eosin (H&E) staining was performed to evaluate histopathology, and RNA-Seq was conducted on tissues from all three regions. H&E staining revealed few effects on the duodenum, but increasingly pronounced disruption and epithelial alterations in the jejunum and ileum, respectively. RNA-Seq analysis of controls compared to the high-dose (HD; 1.5 mg/day) group supported these observations. The number of differentially expressed genes (DEGs) was low (19) in the duodenum, higher in the jejunum (114 DEGs), and significantly greater in the ileum (4982 DEGs) suggesting differential sensitivity of the different regions of the small intestine. Genes associated with immune response, ion transport, and junctional proteins were among the groups of genes most differentially expressed. These results demonstrate the potentially harmful effects of PS NPs on the small intestine of mice and that the sensitivity to NPs increases along the intestine.

## INTRODUCTION

The types, presence, uses, and distribution of plastics has increased exponentially since their introduction in the 1950s (Andrady and Neal 2009; Geyer et al. 2017). In recent years, there have been alarming reports of the accumulation of plastics in the environment, resulting from their ubiquitousness and inadequate waste management (Eriksen et al. 2023). As a result of this, plastic particles have been found throughout the environment and have accumulated in a variety of both aquatic and terrestrial wildlife species (Kaur et al. 2024).

Multiple studies have also reported the presence of plastic particles in humans. Of particular concern are studies which have reported microplastics (MPs) and nanoplastics (NPs) in numerous human tissues, including the male reproductive tract, the placenta, blood, and the brain, among others (Yong et al. 2020). These findings have led to a surge of interest regarding the effects of plastics and their degradation products, namely MPs and NPs. MPs and NPs result from processes such as industrial synthesis, abrasion (eg car tires), washing and wearing of textiles, or from weathering of larger plastic containers, coated surfaces, and fabrics by UV degradation, heat, oxygen, enzymatic degradation and microbial colonization. MPs are generally defined as plastic particles ranging from 5 mm to 1 µm, while NPs are 1µm to 1 nm (Hartmann et al. 2019). In addition to size, MPs and NPs are also categorized according to various other properties, including chemical composition, shape, structure, charge and solubility (Hartmann et al., 2019). As such, it is obvious that assessing environmental and biological impacts of MPs and NPs is complex and difficult to elucidate. Similarly, administration, dose, and route of exposure to MPs or NPs can both complicate and influence observations of any effects on organisms or environmental system. The chemical composition of plastic polymers also varies to include polyethylene, polystyrene, polypropylene, terephthalate, polyamides, polymethyl methacrylate, and polyvinyl chloride (Yin et al. 2021). A study comparing the accumulation of plastic polymers in the testis of dogs and humans living in the same geographical area reported that propylethylene and polyvinylchloride were found to be the most highly accumulated plastic polymers (Hu et al. 2024). In another study examining plastic accumulation in mice and humans from an animal facility, the authors identified five types of MPs including polyamide 66, polyvinylchloride, polyethylene, polystyrene and polymethyl methacrylate (Yang et al. 2023). These studies highlight not only differences in size of plastic particles but also the fact that humans are exposed to a multitude of plastic polymers (He et al. 2023).

It is generally believed that smaller NPs can easily cross epithelial barriers and enter cells. The increased surface of these molecules allows them to interact with organelles, including the mitochondria (Kong et al. 2025). It has been suggested that large molecules are associated with effects at the levels of the plasma membrane, resulting in structural effects and tears in the plasma membrane (He et al. 2023). Several studies have reported the toxicity of MP/NPs in a variety of organ systems including the intestine of both mice and zebrafish (Ghosh et al. 2026; Sun et al. 2024b). Sutton and Hills (Sutton and Hills 2025) suggested that in the intestine there are three stages of NP toxicity: oxidative stress; barrier destruction due to increased inflammation; and microbiome reconfiguration. Reported effects on the intestinal barrier have included a loss in tight junctions due to decreased levels of occludin (OCLN), claudins (CLDNS) and tight junction protein 1 (TJP1 or ZO1), and these were associated with an increase in a variety of cytokines (Chen et al. 2024). In addition, structural changes to the epithelium, including a loss in mucus-secreting goblet cells, were noted and associated with the overall loss of barrier function (Sun et al. 2024a).

Studies have indicated that the accumulation of MPs in the intestine can alter the microbiome and that these MPs can translocate and exert toxicity on other organ systems including the liver, pancreas and brain (Yuvika et al. 2026). The toxicity of these appears to be linked to the plastics’ ability to induce reactive oxygen species (ROS) in target organs, resulting in increased inflammation (Yuvika et al. 2026). In the duodenum, it was reported that MP exposure resulted in increased inflammation leading to increase edema and vacuolization, increased depth of crypts, and alterations in the cilia (Djouina et al. 2022; Liu et al. 2020; Qiao et al. 2019). Studies on the gastric epithelia of the mouse indicate that exposure for 28 days to 50 and 250 nm polystyrene (PS) NPs resulted in alterations to mitochondrial functions and that these resulted from increased ROS production and activation of the p62/Keap1/Nrf2 pathway (Sun et al. 2024b). Cellular effects of MPs using Caco-2 cells also reported effects on the mitochondria (Wu et al. 2019). These studies indicate that the intestine is not only a primary site of NP absorption but also of toxicity. The main route of exposure to MPs and NPs for humans and other biological organisms is from food and their ingestion (Cox et al. 2019). Few studies, however, have focused on long-term exposure to NPs to assess their consequences on the intestine.

The objective of the present study was to observe and determine the effects of a subchronic daily exposure (60 days) of polystyrene (PS) NPs on the mouse small intestine and to compare effects and gene expression in the three regions of the small intestine.

## MATERIALS AND METHODS

### Animals

Six-week-old C57BL/6 male mice were purchased from Charles River Laboratories (Kingston, NY) and acclimated for 1 week under a constant photoperiod of 12h light/dark cycle). Mice received food and water *ad libitum*. All animal protocols used in this study were approved by the university animal care committee.

Following acclimation, mice were weighed and placed randomly into the following groups: control (sterile water); LD (low dose; 0.15 mg NPs/day); HD (high dose; 1.5 mg NPs/day). Doses were selected based on published data regarding environmental levels of NPs and estimates of NP levels in human and animal species. The LD was calculated as being within the range of environmentally relevant NP exposure, while the HD represented a dose just above the upper range (Lu and Chen 2020; Xie et al. 2020). A total of 8 mice were used for each group. Mice were weighed weekly and verified daily for any indication of ill health. Mice were gavaged at approximately the same time every day for 60 days. The following day (day 61), mice were anesthetized with isofluorane and euthanized by cervical dislocation. Tissues were removed and placed immediately in either Bouin’s fixative for immunohistochemistry or frozen in liquid nitrogen for subsequent RNA isolation.

### Nanoplastics

Sterile non-functionalized polystyrene plastic nanospheres (NPs; 500 nm) were purchased from Alpha Nanotech (Vancouver, BC, Canada). Scanning electron microscopy was done for size verification and characterization (diameter = 499.3 ± 5.2; PDI=0.080 ± 0.052; zeta potential(mV)= −55.6 ± 0.6) as previously described (Tavakolpournegari et al. 2026). Test solutions were prepared under sterile conditions and diluted to the desired concentration using sterile filtered H_2_O. The concentration of the solution was determined such that a volume of 0.2 ml delivered the equivalent of either 0.15 or 1.5 mg NP per mouse per day.

### Histology

Following euthanasia of the mice, tissues designated for histology were rinsed briefly in PBS, blotted dry, subdivided into the duodenum, jejunum and ileum and fixed in Bouin’s solution. The following day, tissues were removed from Bouin’s, rinsed in 70% ethanol to remove excess fixative, and stored in fresh 70% ethanol until tissue processing. Tissues were dehydrated using graded ethanol washes, xylene substitute/ethanol and xylene substitute prior to embedding into paraffin. Tissues were sectioned (6 µm) and mounted on a glass slide by either the Goodman Cancer Research Center (McGill University, Montreal, QC) or l’Institut de Recherche en Immunologie et Cancerologie (Université de Montréal, Montreal, QC).

### Immunohistochemistry

Sections used for immunohistochemistry were rehydrated in 3 changes of xylene substitute (MilliporeSigma Canada, Oakville ON) and then through a series of graded alcohols, including a saturated lithium carbonate alcohol (70 % EtOH) bath, to neutralize any picric acid from Bouin’s fixation. An additional 3% hydrogen peroxide bath was performed to quench any endogenous peroxidase activity. Sections were then subjected to heat-mediated antigen retrieval, using citrate buffer (pH 6.0), heated at 70% power in a microwave oven for 10 minutes, and allowed to cool to room temperature prior to subsequent steps. The sections were then rinsed 3 times in H_2_O and once in Tris-buffered-saline with 0.1% Tween-20 (TBST), prior to blocking for 1 hr at room temperature in TBST plus either 5% serum of host secondary antibody species or 5% bovine serum albumin (BSA; BioBasics, Markham ON). The appropriate primary antibody (Suppl Table 1) was then diluted in blocking solution and tissue sections were incubated for 2 hrs at room temperature or overnight (18 hrs) at 4°C in a humidified chamber. Sections were subsequently washed 3 times in TBST and then incubated for 1 hr with the appropriate horseradish peroxidase-conjugated secondary antibody (Invitrogen, Burlington, ON; Supplementary Table 1). Sections were washed 3 times in TBST, dehydrated through graded alcohols and xylene substitute, and mounted with Permount (Fisher Scientific).

### Nucleic acid extraction

Frozen tissues were weighed and then ground under liquid nitrogen, using sterile mortar and pestles, and placed immediately into sterile tubes containing lysis buffer (NucleoSpin RNAPlus kit, Macherey-Nagel, Mississauga ON). The resulting lysis buffer-tissue suspension was then frozen in liquid nitrogen and stored at −80C until further processing.

RNA was extracted according to manufacturer’s instructions using the NucleoSpin RNAPlus kit. The quality and concentration of the resulting RNA was verified using a Nanodrop apparatus (Fisher Scientific), and subsequently treated with DNase (Ambion, Mississauga, ON) to remove any residual DNA. Quality and concentration of the final RNA was again verified using the Nanodrop and RNA was immediately frozen in liquid nitrogen and stored at −80C until further use.

#### RNA Sequencing (RNAseq)

Total RNA isolated from tissues of each of the three regions of the small intestine (duodenum, jejunum, ileum) from control (water) and treated (1.5 mg NP/day) groups were sent to Genome Quebec (Montreal, QC) for sequencing. The quality of isolated RNA was assessed using an Agilent 2100 Bioanalyzer (St. Laurent, QC). Samples with RNA integrity number (RIN) ≥ 7.0 were used for RNASeq library preparation following poly(A) selection and sequencing performed by standard Illumina platform protocols (NovaSeq PE 100 – 25M reads; Genome Quebec). The resulting raw sequences were deposited in the NCBI Gene Expression Omnibus (GEO) repository under the accession number GSE (pending).

The raw sequencing reads were processed by the Bioinformatics core facility of the Montreal Clinical Research Institute (IRCM). The quality of raw reads was assessed using FASTQC software (v0.11.8) followed by trimming with TRIMMOMATIC software (v0.36). The reads were aligned to the mouse reference genome GRCm38 using STAR software (v2.7.6a) and the raw counts were quantified with FeatureCounts (v1.6.0). Bioinformatic analysis for differential expression analysis was performed in R using the DESeq2 package (v1.52.0) which generates differential expression results with p-values adjusted using the Benjamini–Hochberg procedure to control the false discovery rate (FDR). The DEG identification and the subsequent analyses were based on these adjusted p-values (Love et al., 2014). Samples were categorized into control and treatment groups for pairwise comparisons. Significantly differentially expressed genes (DEGs) were those with an adjusted p-value < 0.05. Principal component analysis (PCA) plots were generated to explore the overall expression patterns and sample clustering for each intestine region. Volcano plots based on p-value and shrunken log2FoldChange (log2FC) were generated from the DESeq2 results to view comparisons between control and treated mice (Zhu et al. 2019). Additionally, the top 25 DEGs (based on p-value) from each comparison were visualized using heatmaps to highlight the group-specific expression profiles. For visualization purposes due to the large number of DEGs in the ileum log2FC was used as cutoff setting at 8 for upregulated genes and −8 for downregulating genes. Heatmaps were also used for representing clustering based on families of genes, such as immunoglobulin production, endocrine responses, cadherins, tight junctions and defensins.

Protein class identification of highly regulated gene was done using the PANTHER Knowledgebase (Mi et al. 2021). DEGs were also subjected to STRING Network analysis v11.0 (Szklarczyk et al. 2023) to assess relationships between proteins as a network.

### qPCR

Purified RNA of control (water) and HD NP groups was reverse-transcribed (RT) to produce cDNA using the LunaScript RT Supermix kit (New England Biolabs) according to the manufacturer’s instructions. Following RT, the samples were then added to a reagent mixture containing Luna universal Mastermix (New England Biolabs), specific forward and reverse primers, and sterile water, in a total reaction volume of 20 µl (per sample). All samples were done in triplicate. These samples were then subjected to qPCR in a Corbett Rotor Gene 3000 thermocycler for 20-40 cycles, depending upon the gene of interest (Supplementary Table 2). Each run included a sample for normalization and samples of serially diluted standards. Data were normalized to control or housekeeping genes using the two standard curve method (Livak and Schmittgen 2001). Student T-tests and One-way ANOVA were used to assess statistical significance, defined as \**p* ≤ 0.05 and \*\**p* ≤ 0.01.

#### Transepithelial Resistance in Caco-2 Cells

Caco-2 cells (ATCC-HTB-37) were grown according to standard culture protocols on T75 Cell+ flasks (Sarstedt, Germany) and maintained for several passages prior to commencement of experiments. Cells were cultured in EMEM medium supplemented with 10% Fetal Bovine Serum (FBS), 1% penicillin-streptomycin, 1% MEM non-essential amino acids, 1% HEPES, and 1 mM sodium pyruvate (Wisent, St-Jean-Baptiste, QC). Cells were then seeded on Corning Transwell inserts (0.4 µm pore size; Fisher Scientific) for 24-well plates, at a density of 20,000 cells/cm^2^. Sterile PS-NPs (500 nm, Alpha Nanotech, Vancouver Canada) were diluted into medium at 4 different concentrations: 0 (controls), 25, 50, and 100 µg/ml. Transepithelial electrical resistance (TEER) was measured with an EVOM 2 instrument (Epithelial Volt/Ohm Meter, World Precision Instruments, Sarasota FL) at 24, 48, 72, and 96 hrs following addition of PS-NPs. Triplicate wells for each condition were prepared. Data are presented as the mean ± SEM from independent biological replicates for *in vitro* experiments (n=3). Data were analyzed using two-way ANOVA followed by Dunnett’s test. Significance was set at p<0.05.

## RESULTS

Mice exposed to either vehicle or low and high dose of PS-NP for 60 days did not display any obvious behavioral changes or differences in body weight at the end of the study. Tissue sections from each of the mice in all experimental (NP-treated) and control groups were stained according to standard hematoxylin/eosin protocols. Examination of these sections revealed normal morphology of most cells of the epithelium in the duodenum, jejunum, and ileum. However, as seen in Fig. 1, epithelial aberrations were observed in all regions of the intestine in mice treated with both low (LD; 0.15 mg/day) and high (HD; 1.5 mg/day) groups. These morphological changes included increased vacuolization and morphological disruption of the epithelium in some areas (Fig 1). These vacuoles were present throughout the intestine and their frequency increased from proximal (duodenum) to distal (ileum) regions of the small intestine. While some vacuoles were small the presence of larger vacuoles were observed in the jejunum and ileum (Fig 1). In the ileum there was extensive tissue damage observed in mice treated with both the low dose and high dose of PS-NP. The epithelium appeared disorganized, indicative of substantial alterations in this region of the intestine.

**Figure 1.**
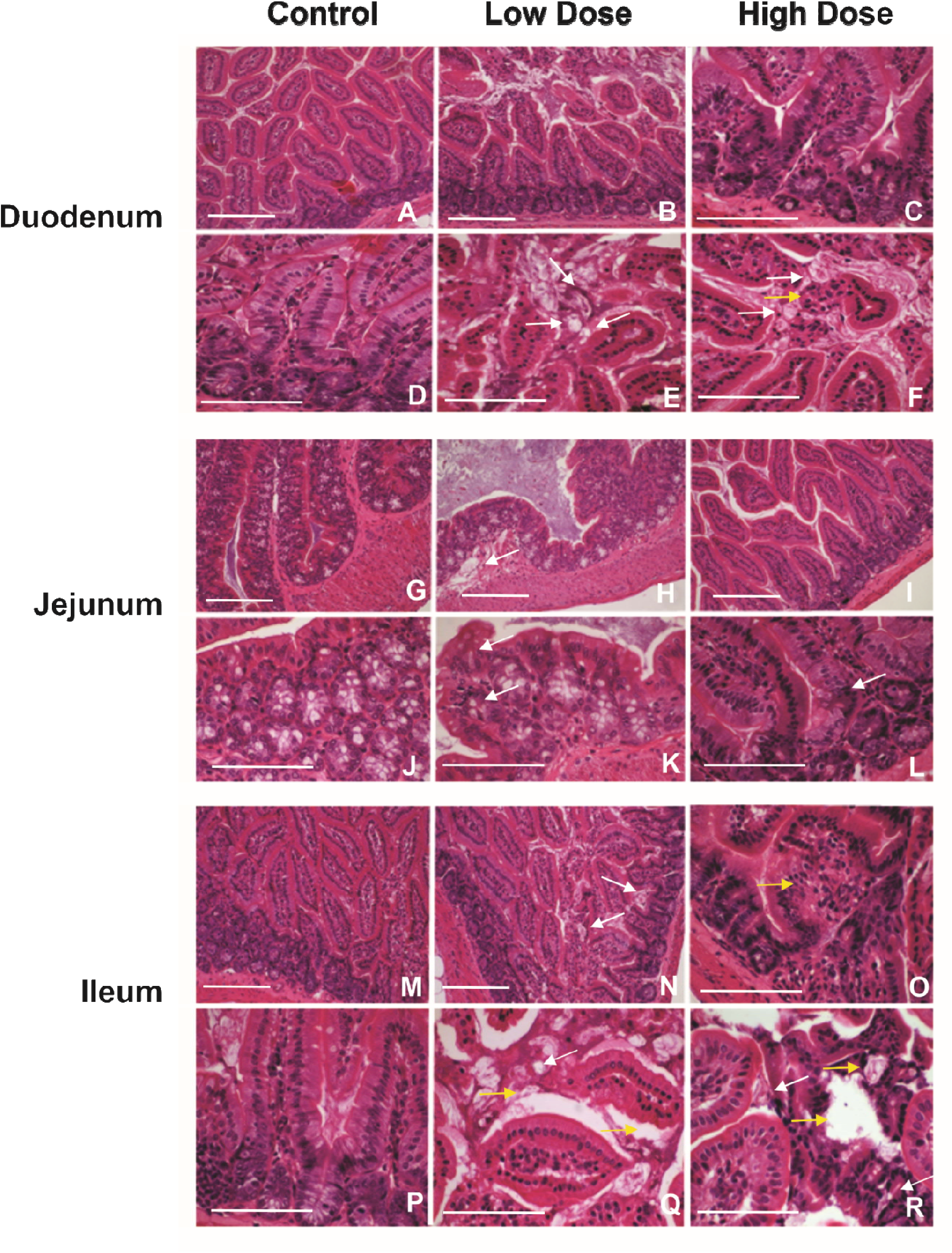
Tissue sections from the 3 regions of the intestine of mice exposed for 60 days to water (vehicle control); low dose, (0.15mg/day) and high dose (1.5 mg/day) of PS-NP by gavage (n=8 mice per experimental group). Sections were fixed in Bouin’s solution, embedded in paraffin and stained with H&E. Representative images are shown for each region of the intestine. In the duodenum (A-F) and in the jejunum (G-L) small vesicles were apparent (white arrows). Tissue abnormalities including hyperplasia in duodenum were also noted in the high dose group (yellow arrow). In the ileum (M-R) of mice exposed to either the low dose or the high dose, there were notable abnormalities in the epithelium and detachment of the epithelium from the basement membrane (yellow arrows). Scale bar = 50µm

To elucidate the cause(s) of morphological changes noted in the intestinal epithelia of NP-treated mice, RNAseq analysis was done comparing controls and high dose exposed mice for each of the three regions of the small intestine (duodenum, jejunum and ileum). Principal components analysis indicated that RNAseq from both the jejunum and ileum were notably different between controls and treated mice. However, there was greater overlap between controls and treated mice in duodenum (Fig 2). Volcano plots indicated that differentially expressed genes (DEGs) varied in the different regions of the intestine (Fig 3). Data indicated that there were relatively few (19) DEGs in the duodenums of these mice. (Fig 3). Among the 19 DEGs, 7 genes were up regulated and 12 genes were downregulated. Although this was initially surprising, these data correspond to the somewhat less extensive morphological changes observed in the epithelia of PS-NP exposed mice, as evaluated by H and E staining (Fig 1). In contrast, there were 114 DEGs in the jejunum of PS-NP-exposed mice versus control. (Fig 3). These included 22 DEGs that were decreased and 92 DEGs that were increased, including 4 DEGs that were decreased and 18 DEGs that were increased by at least 10-fold. In the ileum, there was a substantial increase in the number DEGs with 4982 significantly different DEGs. Of those, 2733 genes were increased and 2249 genes were significantly decreased. A change of 10-fold or more was observed for 365 genes which increased and 269 genes which decreased, showing a substantial effect on almost 650 genes in the ileum. These data reflect the dramatic changes observed in the morphology of the ileum in PS-NP treated mice (Fig 1). RNAseq data was validated by qPCR which showed similar results to those obtained by RNAseq (Suppl Fig 1).

**Figure 2.**
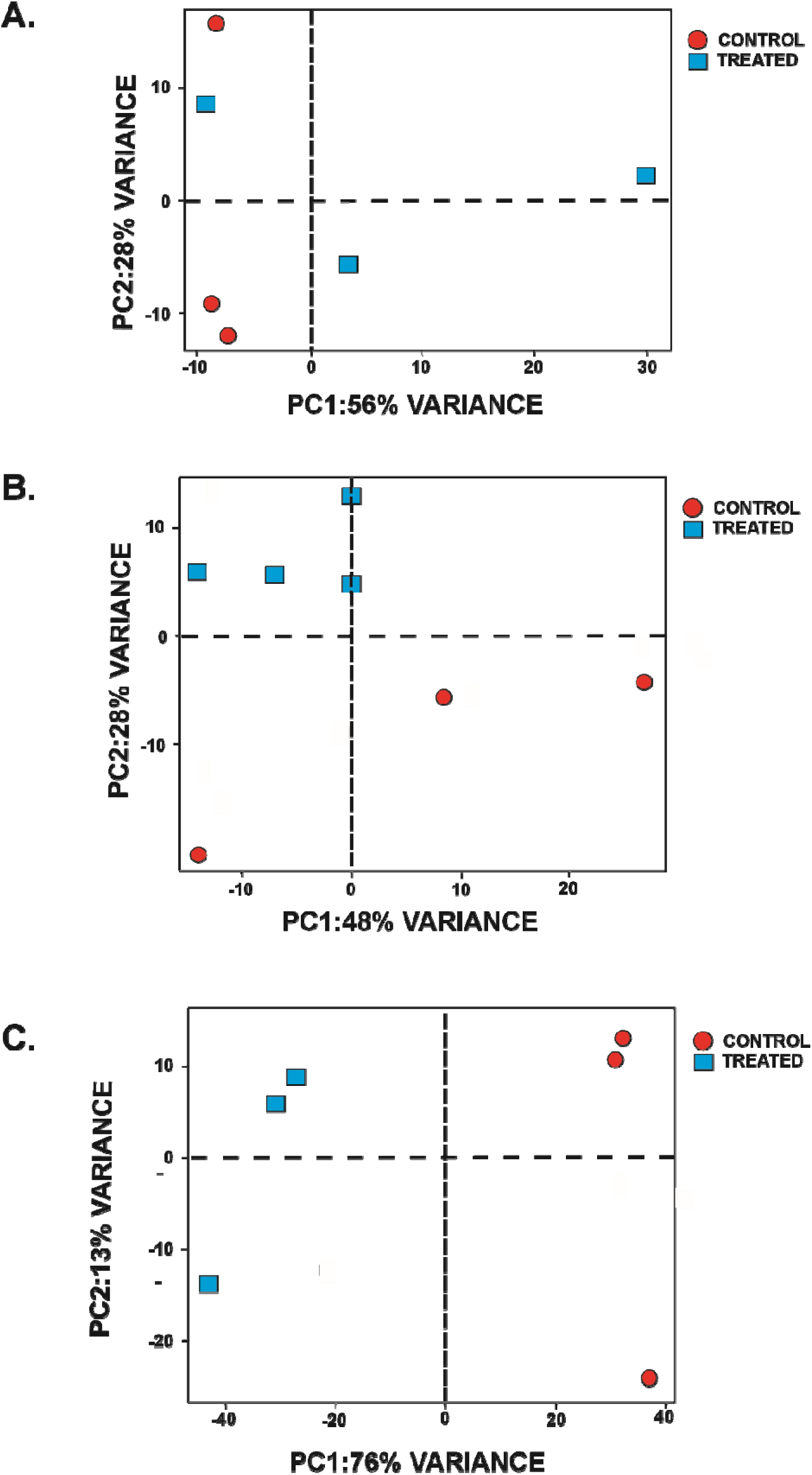
Principal component analysis of summarized RNA Seq data for control and high dose PS-NP exposed mice for 60 days. Graphs show differences between control and treated mice for each of the three regions of the intestine.

**Figure 3.**
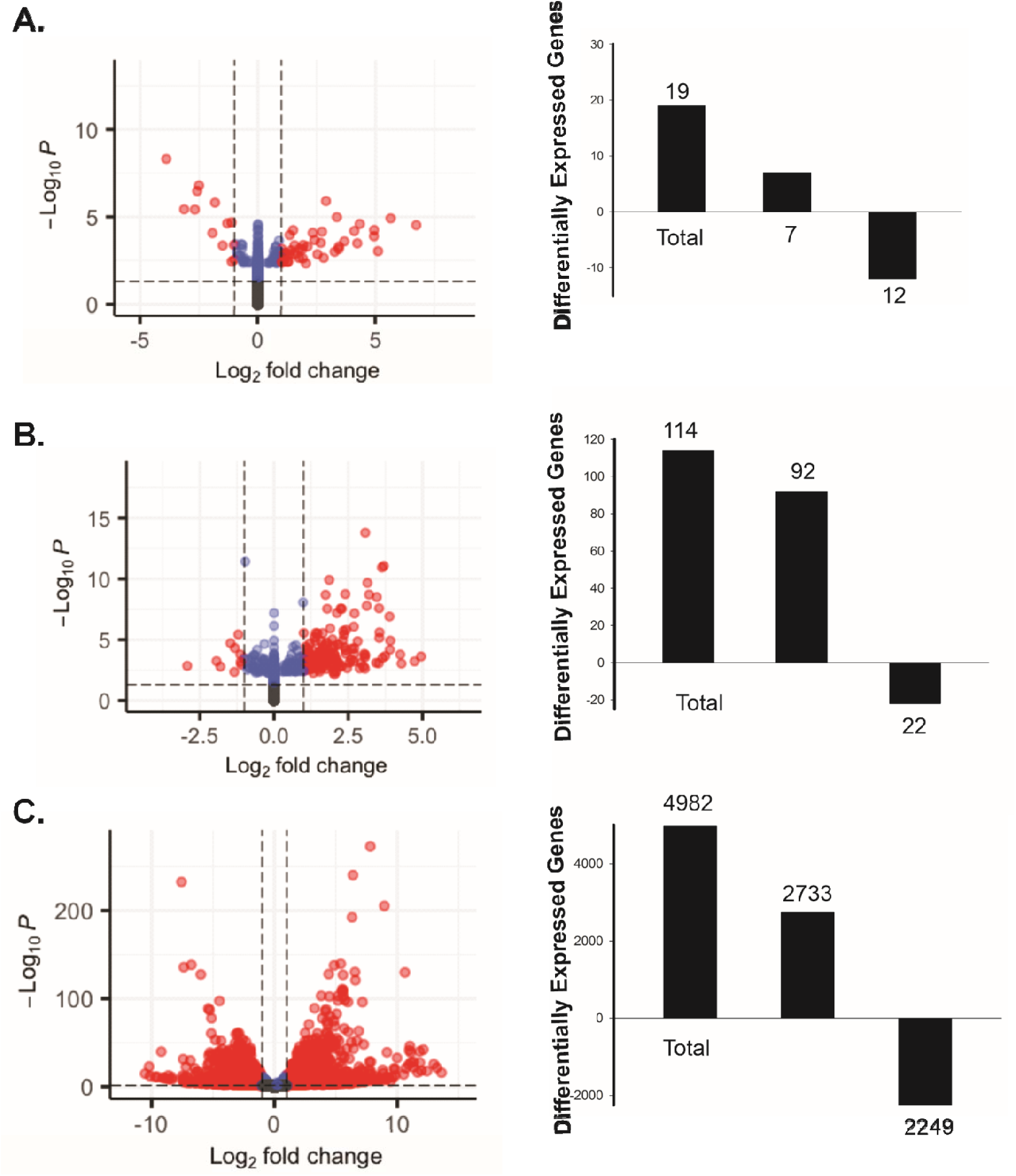
Volcano plots displaying RNA Seq data from high dose (1.5 mg/day) versus control groups for each of the regions of intestine aligned across (A-duodenum, B-jejunum and C-ileum). The number of differentially expressed genes (DEGs) is shown in histograms displaying the total number of DEGs, as well the number of significantly increased and decreased genes. Note the high number of DEGs in the ileum of treated mice.

Heat maps of DEGs in the duodenum, jejunum and ileum showed clear differences in gene expression (Fig 4). Some of the genes in the duodenum indicated minor differences between one of the treated mice, which displayed a more distinctive decrease in gene expression. Interestingly, the duodenum displayed a decrease in multiple genes which code for immunoglobulin related genes. In contrast, in both the jejunum and the ileum there was a substantial increase in genes implicated in immune function. Various immunoglobulins and regulators of adaptive immunity were increased in the jejunum (Fig 5). Most of the highly DEGs were those that coded for immunoglobulins (Suppl Table 3). Among the most highly decreased DEGs in the ileum were genes implicated in various aspects of cellular metabolism, cellular transport and vitamin D binding (Suppl Table 3). A substantial number of genes, primarily among the top 20 DEGs in the ileum, coded for defensins (Fig 5; Suppl Table 4). Interestingly, multiple genes implicated in thyroid hormone metabolism were also decreased in the ileum, in addition to dysregulation of both tight and adherent junction proteins (Fig 5). Furthermore, there were no consistent effects on the inflammatory response in the jejunum (not shown). In contrast, several genes were altered in the ileum of HD-exposed mice (Fig. 5). Notable changes included the pro-inflammatory cytokines including IL1 (α and β), TNF, CXCL1, CCK3, CCL4, and Nfκb1as well as anti-inflammatory genes such as IL10, and TGFβ2 (Fig 5).

**Figure 4.**
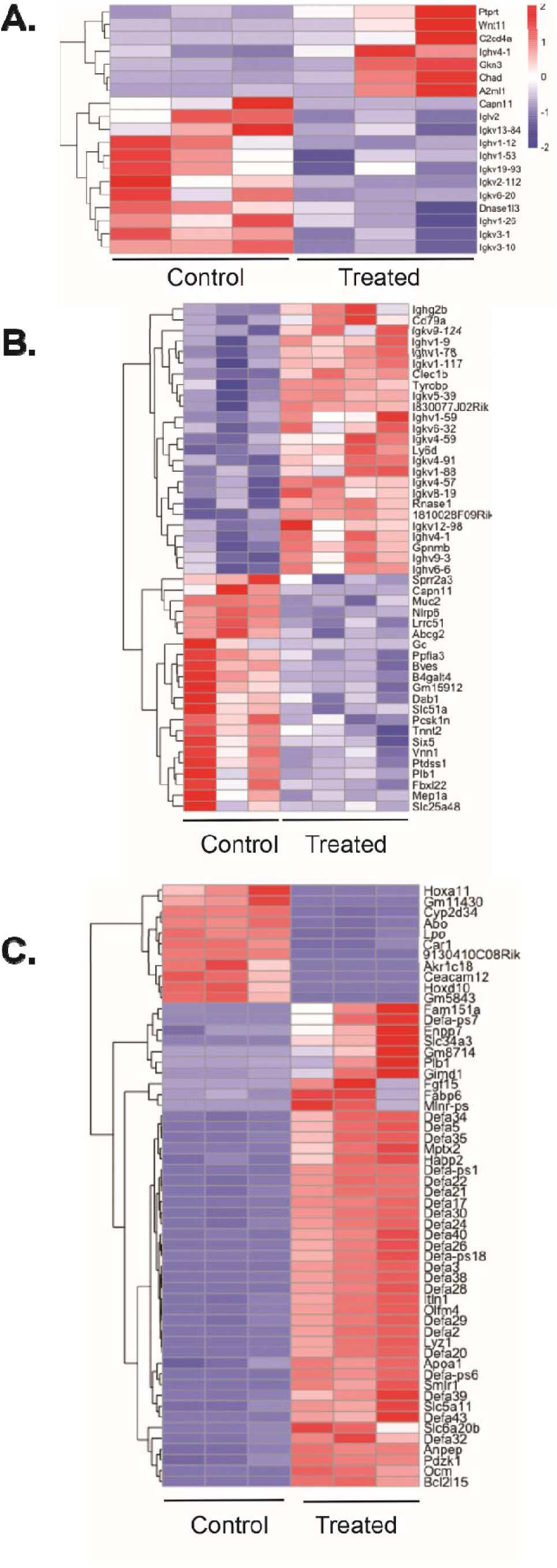
Heat maps of RNA Seq data from high dose (1.5 mg/day) versus control groups for each of the regions of intestine. Note the much higher number of differentially expressed genes (DEGs) in the ileum, as compared to the much lower number of DEGs in the duodenum and jejunum regions. The data clearly indicate a marked difference in gene expression between down regulated (blue) and upregulated genes (red).

**Figure 5.**
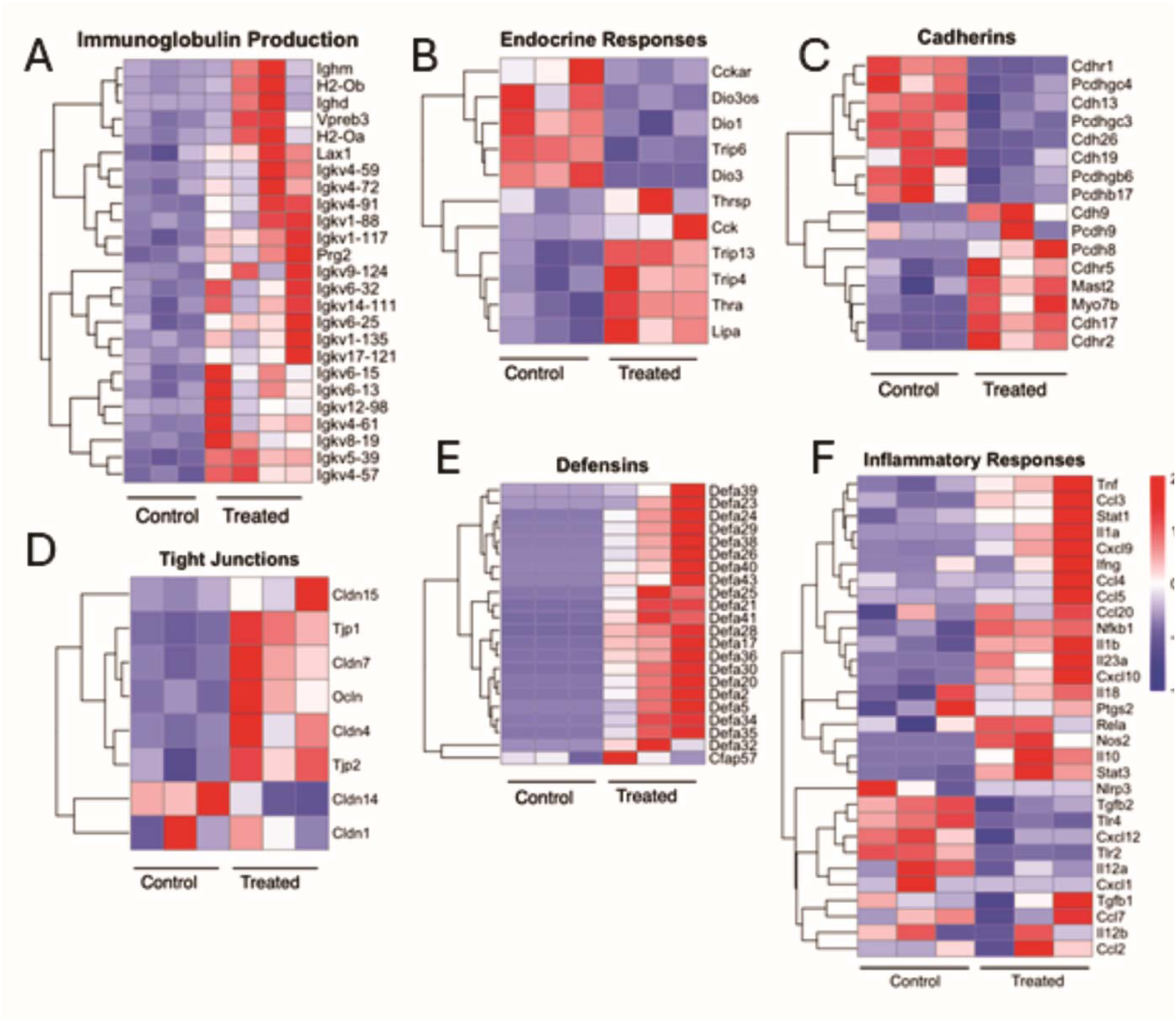
Heatmaps of groups of genes differentially expressed in the jejunum (A) and the ileum (B-F) of control and high dose PS-NP exposed mice. Genes implicated in the production of immunoglobulin (A) were globally increased in the jejunum part of the intestine of the NP treated mice. Differential expression of genes implicated in endocrine responses (B) is shown for the ileum, as well as genes encoding components of adherens junctions (cadherins; C) and tight junctions (D). Expression levels of genes of the defensin family (E) and inflammatory responses (F) were increased in the ileum of NP exposed mice compared to controls.

Grouping of genes by protein class in each of the three regions of the intestine using the PANTHER software indicated that most transcripts in the duodenum showing an increased expression were associated primarily with defence/immunity, protein modifying enzymes, intracellular and transcellular signaling (Fig 6A). In the jejunum, genes related to protein binding were the largest category of transcript functions, followed by catalytic activity, and molecular transducers (Fig 6A). Finally, in the ileum, catalytic conversion enzymes, protein modifying enzymes and cellular transporters were the three largest categories of differentially expressed genes.

**Figure 6.**
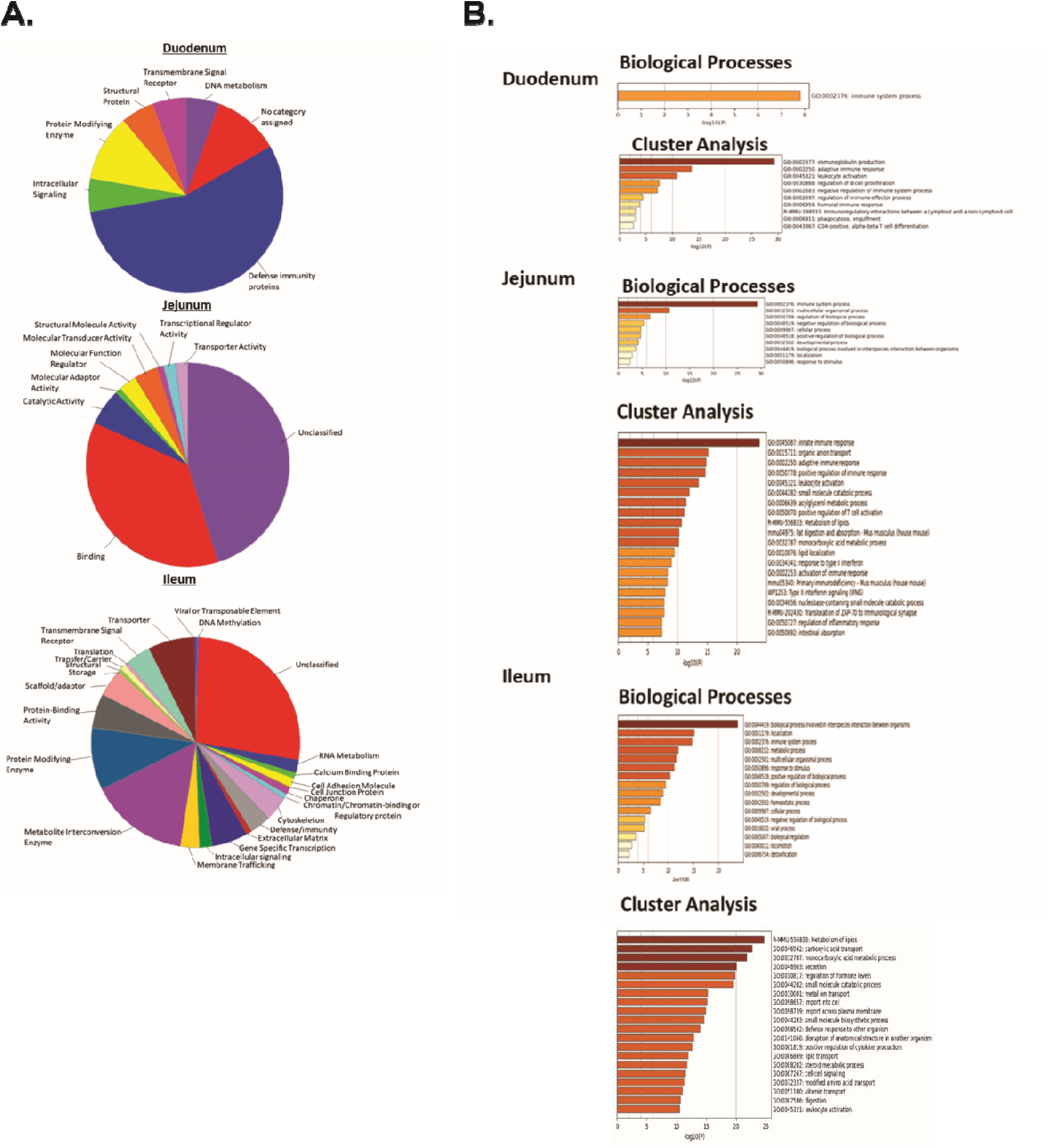
DEGs of each intestinal region grouped according to protein classes, as defined by Panther database (A). In the duodenum, defensin/immunity genes were most differentially expressed, while genes for binding proteins represented the largest group of DEGs in the jejunum. In the ileum, genes related to metabolic enzymes comprised the largest group of DEGs. RNA-Seq data analyzed using GO analysis in the Metascape software indicated that immune function was noted throughout the intestine for both biological processes and cluster analysis.

Enrichment of gene ontology (GO) analysis with Kegg annotations using the Metascape software (Zhou et al. 2019) to examine Biological Processes indicted that all three regions of the intestine showed altered aspects of immune function. Furthermore, in both the jejunum and ileum regulation of cellular, developmental and biological processes were also significantly altered by exposure to PS-NPs (Fig 6B). Cluster analysis also supported the fact that several aspects of immune function were significantly altered in the duodenum, jejunum and ileum. Furthermore, in the ileum, cluster analysis also predicted effects on lipid metabolism, carboxylic transport, monocarboxylic acid metabolism, secretion, hormone regulation and effects on multiple transporters (Fig 6B).

String network analysis was done to identify relationships between proteins as an overall network. String network analysis was not possible for the duodenum due to small number of genes whose expression was altered by PS-NP treatment (Fig 7). In the jejunum, several networks were predicted to interact with one another. Some of the nodes within the network were identified and appeared to point to interesting network interactions including genes involved in immunoglobulins and B-Cell function (Pou2af, Cd79a, Tyrobp, Fcer1) and Was, which is implicated in an actin filament reorganization (Fig 7a). In the ileum, the network was far more extensive due to the greater number of genes that were altered by treatment. String network analysis predicted several developmental signaling pathways in which key nodes were predicted. These included Stat1, Kak2, Plcg2, Rhoa, Wnt3 and Ctnb1. Interestingly, several histone genes were identified. Nodes containing Zap70 and Cd8a, which is implicated in immune regulation were also observed (Fig 7b). Together, the String network shows that cellular differentiation and immune function may explain, at least in part, more extensive effect associated with PS-NP exposure.

**Figure 7.**
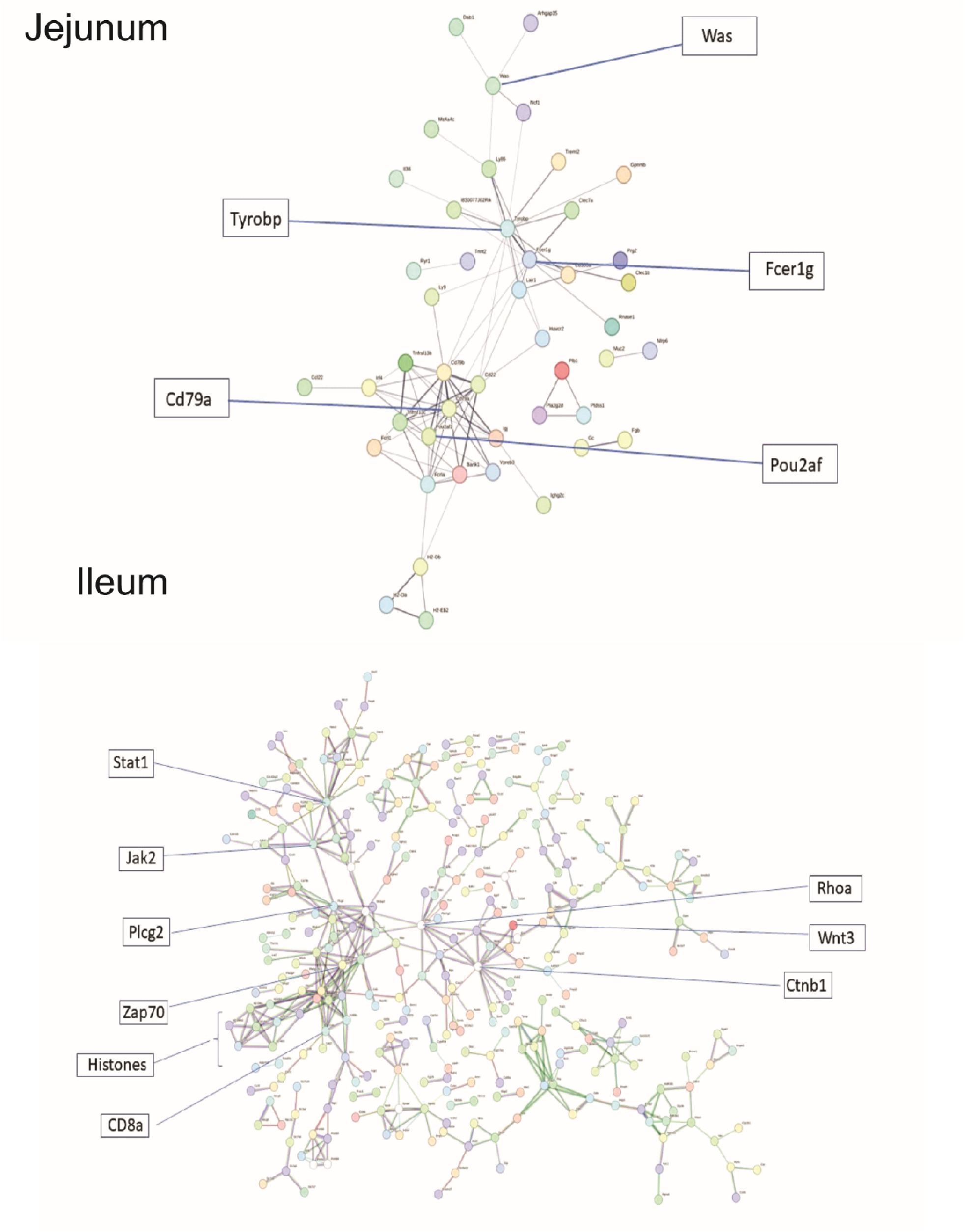
RNA-Seq data as analyzed using the STRING network analysis software, showing potential protein interactions and signaling pathways associated with the DEG data. Note that there were no interactions observed for the duodenum. Several nodes were identified as potentially important links, useful for furthering overall understanding in predicted String Network analysis for both the jejunum and Ileum (squares).

To further explore the morphological alterations observed in histopathology and RNAseq data immunohistochemistry (IHC) of select genes was performed. The target proteins included the adherens junction protein Cadherin 1 (CDH1), and Musashi-1. Musashi-1, a stem cell marker in the intestine, was chosen to see whether any indicators of epithelial renewal were changed. IHC results for Musashi-1 suggest that levels of this protein were reduced in both treatment groups (0.15 and 1.5 mg PS-NPs/day) as compared to controls (Fig 8). CDH1 appeared to display comparable localization of immunostaining among controls and the low-dose (0.15 mg PS-NPs/day) groups in all regions of the intestine (duodenum, jejunum, ileum). For both controls and the low-dose groups, the immunostaining for CDH1 appeared to be more intense in the distal portion (ileum) of the small intestine as compared to the proximal region (duodenum) (Fig 8).

**Figure 8.**
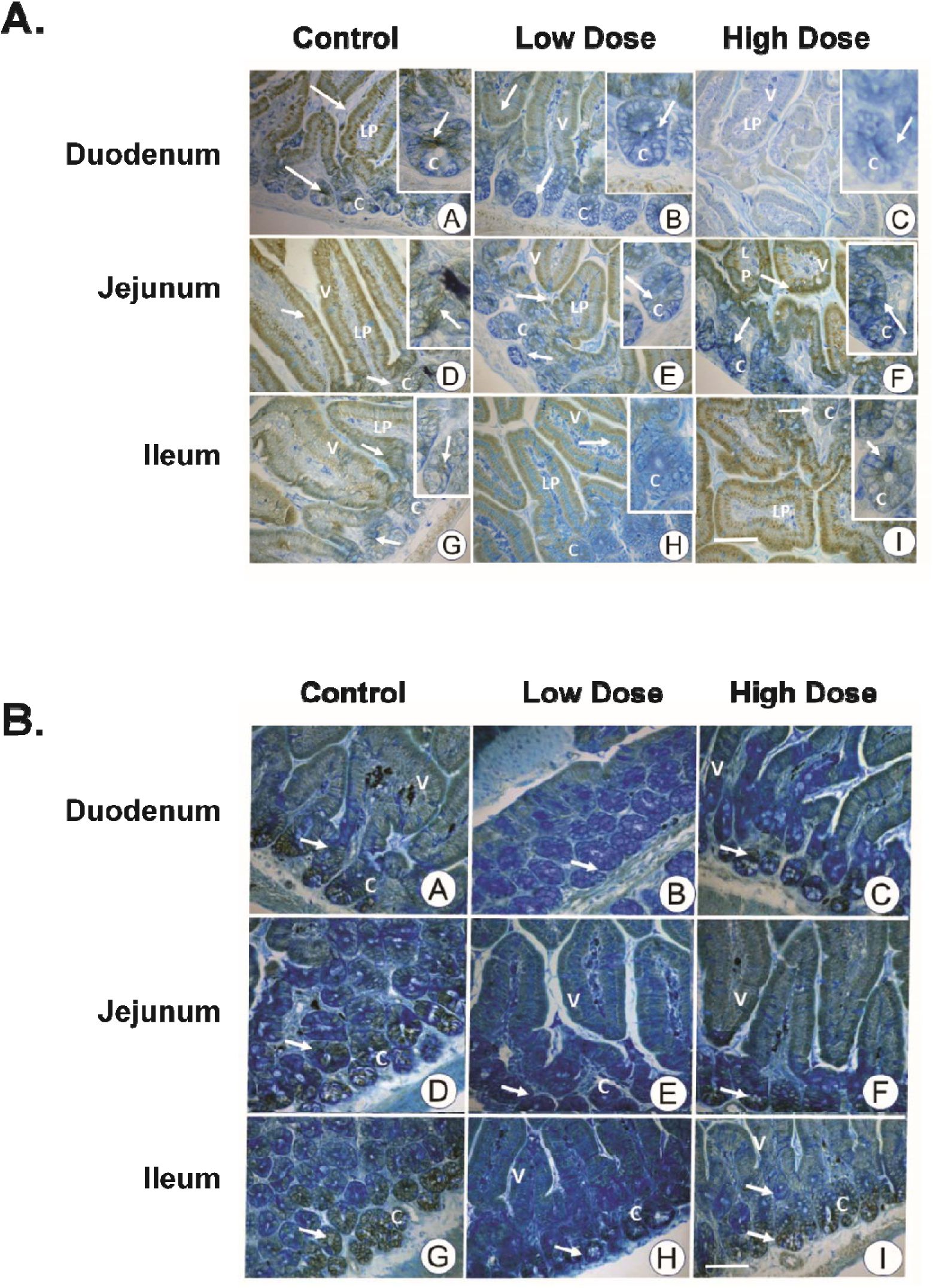
IHC Images of tissue sections from each of the 3 intestinal regions immunostained for either CDH1 (A) or Musashi1 (B). Arrows indicate immunostaining. CDH1 immunostaining was markedly decreased in both the low and high dose groups. There were no obvious distinctions noted for Musashi1 immunostaining. Abbreviations: V-villi; LP-lamina propria, C-crypts. Scale bar = 50µm.

To further examine the effects of PS-NPs on tight junctions, Caco-2 cells were exposed to: 0 (controls), 25, 50, and 100 µg/ml for up to 72 hrs. There were small but statistically significant decreases in TEER for the 100 µg/ml group at 24, 48, and 72 hrs (Fig 9). There were no significant differences at 96 hrs. A trend towards a slight decrease in TEER was noted in the 25 and 50 µg/ml groups at these time points, but the differences between these conditions and the controls were not significant. A decrease in TEER in PS-NP-treated cells would suggest a disruptive effect on the junctions between these cells.

**Figure 9.**
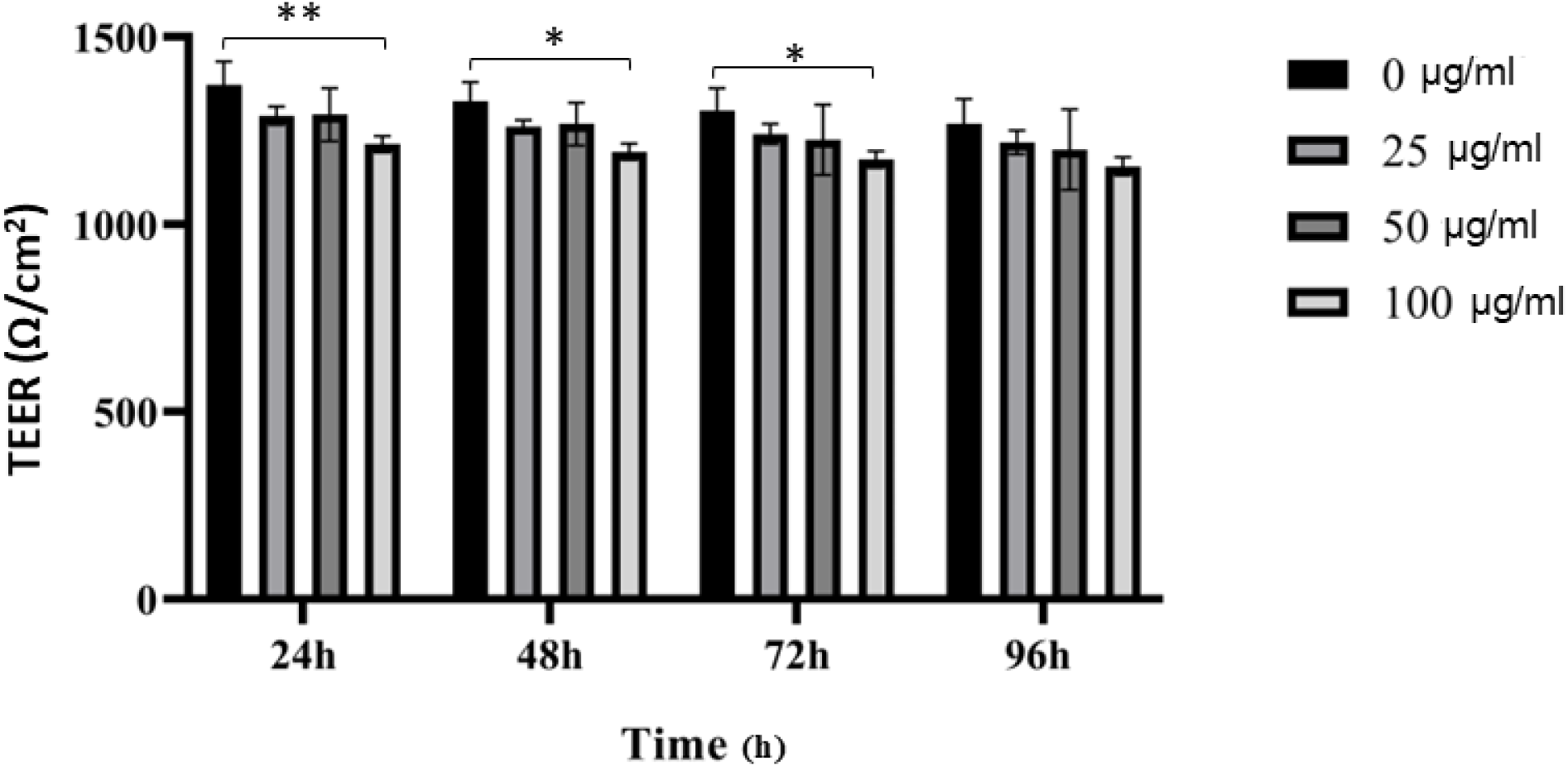
TEER in Caco2 cells exposed for up to 96 hrs to 0 (controls), 25, 50, and 100 µg/ml of PS-NPs. TEER was measured using an EVOM 2 Epithelial Volt/Ohm Meter. All experimental conditions were done in triplicate (n=3). * indicates a significant difference (p<0.05).

## DISCUSSION

There has been a surge of interest in recent years regarding the uses, management and waste disposal of MP and NP, as well as the increased presence of NPs in air, water, and soil environments, and in organisms living in these environments. Ingestion is the most common route of NP exposure in organisms. The current study reports the results of a sub-chronic (60 days) oral exposure (gavage) of polystyrene nanoplastics (NPs) to mice. The intent was to provide an initial assessment of long-term PS-NP ingestion to model human exposure, given that ingestion is a major route of exposure, and PS is a widely used plastic.

NP uptake has been examined in several studies, which report 2 main ways for cellular uptake of NPs. These include endocytosis and clathrin-mediated endocytosis (Huang et al. 2023). Studies have shown bioaccumulation in several tissues, including the intestines (Ding et al. 2021; Liang et al. 2021). Others have reported the ability of NPs to cross the intestinal barrier (Okano et al. 2025) and infiltrate intestinal cells (Domenech et al. 2021; Fan et al. 2016). There are reports of cytotoxic effects following NP exposure, as documented in a recent review (Hirt and Body-Malapel 2020). These include ROS production, mitochondrial damage, endocytic system perturbation, decreased cell viability, apoptosis, pro-inflammatory response(Chen et al. 2024; Domenech et al. 2021; Magri et al. 2018; Mahler et al. 2012).

However, other studies have reported no effects following administration or exposure of PS-NPs (Paget et al. 2015; Schirinzi et al. 2017; Walczak et al. 2015). A recent study indicated that intestinal damage from acute colitis was reduced in mice orally exposed to polyethylene terephthalate (PET) MPs and NPs prior to induction of colitis (Okano et al. 2025). Furthermore, these MPs and NPs were generated from actual plastic products, more accurately reflecting environmental relevance. After 5 days of colitis induction, the MP and NP treatment downregulated 138 genes and upregulated 76 genes, many of which were immune-related, consistent with our observations. The number of DEGS was markedly reduced after 8 days, when less than 10 genes were up-or down-regulated. A similar pattern of changes was also seen with respect to genes related to cytokine and chemokine gene expression. The authors suggested that while this modulation of immune function may initially seem good, it may have a negative impact on long-term immune function, as well as an impact on the mucosal intestinal barrier. Mice exposed to PS-NP for 60 days in our study did not exhibit any overt signs of toxicity with respect to body weights or behavior. Epithelia of the three regions of the small intestine indicated morphological changes in mice treated with either a low dose (0.15 mg/day) or high dose (1.5 mg/day) of PS-NPs for 60 days. These changes included increased vacuolization of the epithelium in NP-treated mice, as well as detachment of portions of the epithelium of the ileum at both PS-NPs exposure doses.

Such changes suggested that NPs exerted effects on the intestinal epithelium. Similar epithelial disruptions or damage have also been reported in other studies with PS -NPs in mice for shorter times of exposure(Jin et al. 2019; Liu et al. 2020; Su et al. 2024; Zhang et al. 2024). Conversely, Xiao et al (Xiao et al. 2022) reported no effects on intestinal epithelia in mice exposed for 30 days via gavage to varying concentrations of PS-NPs, as evaluated by H&E sections, inflammatory or reactive oxygen species (ROS) markers. Several mechanisms have been proposed for observed epithelial alterations; the majority of these, attribute any changes to being the result of pro-inflammatory processes or ROS production (Yin et al. 2021; Zhang et al. 2024). It is evident that further research is required to elucidate the mechanisms responsible for the observed epithelial alterations.

Changes in DEGs along the intestine of NP-PS treated mice indicate an increase in changes in overall gene expression from the duodenum to the jejunum to the ileum. All regions of the intestine show alterations in immune function suggesting that PS-NP exposure exert effects on different aspects of immune function. Analysis using GO annotation of biological processes and String network analysis also support the notion that sub-chronic exposure to PS-NP resulted in immune disfunction throughout the intestine. The ileum displayed a vastly greater number (4982) of DEGs in the HD group as compared to either the other regions of the intestine or controls. The data from our experiments show that the ileum was the most sensitive region of the small intestine to PS-NP exposure. This has been observed in other studies as well, with various types and sizes of NPs (Choi et al. 2024; Du et al. 2024; Zhang 2023). Although this effect may point to differences in sensitivity to PS-NP, it does not negate the possibility that PS-NP may accumulate in the region of the intestine after 60 days of exposure. Nevertheless, it suggests that the ileum is particularly sensitive to exposure to PS-NP, resulting is dramatic changes in gene expression.

In our study, MPTX2 was one of the most upregulated genes in the ileum of HD animals. This was corroborated by the qPCR data, which also showed upregulation of this gene in the HD group. Furthermore, there were multiple genes coding for defensins that were upregulated in the Ileum of PS-NP exposed mice. In fact, 15 of the 20 most highly expressed DEGs in this region were defensins. This is consistent with reports of upregulation of these genes in response to infection, inflammation, or chemical exposure (Bashir et al. 2022; Zhang et al. 2023), increased ROS, and antibacterial defense (Yan et al. 2024). Given that changes in ROS have been reported in cells exposed to PS-NPs, this response may be the results of induction by ROS in the ileum (Chakraborty et al. 2026; Das 2023).

A recent review of over 50 articles examined NPs, particularly PS NPs, in which the authors concluded that there were 3 main steps common to NP toxicity on the intestine (Sutton and Hills 2025). It appears that in most situations, ROS are generated by NPs, which can also decrease antioxidant defenses. This results in oxidative injury to both the tissue itself as well as to the microbiota of the intestine, thereby negatively impacting mucus production but activating inflammation. This in turn shifts the microbiota composition to favor stress-tolerant bacterial species, disruption of normal intestinal function, altered immune function, and compromised nutrient absorption and metabolism. More importantly, the authors of this review noted that only PET (polyethylene terephthalate) at low environmentally relevant doses did not trigger the sequence of oxidative stress, disruption of the intestinal epithelial barrier, and altered microbiome. This suggested that for most of the plastic types and particle sizes included in these studies, toxicity could be induced at low doses or concentrations of plastics (Hirt and Body-Malapel 2020; Huang et al. 2023; Sutton and Hills 2025). In contrast, Okano et al (Okano et al. 2025) studied MPs and NPs produced from PET and examined their biological impact on the intestines of mice. They found that in mice exposed first to PETs and then induced with colitis, the degree of intestinal damage was reduced, as compared to those just induced with colitis. Furthermore, the mice exposed to PET and then subjected to colitis displayed a reduced degree of disease. Clearly the role of the microbiome is complex, as are effects of NPs, which may act directly on the intestinal epithelium as well as indirectly via the microbiome. Interestingly, it is evident that in our experiments as well as in other studies, the ileum was the region of the small intestine that was most sensitive to PS-NP exposure.

Additionally, most of these studies also report alterations to various parameters of the immune system functionality, also observed in our study. Although these effects are diverse, given that multiple studies in both vertebrates and invertebrates have examined NP and MP effects on the immune systems, there are commonalities in terms of observed changes in activity of immune-related enzymes, immune cells and cytokines. Nonetheless, no clear pattern of immunotoxicity has emerged, likely due to the many variables regarding type, chemical composition, size, shape, dosages, and routes of exposure, highlighting the complexity of assessing any type of effects due to MP or NP exposure.

*In vitro* studies with the intestinal cell line CaCO-2 cells using PS-NPs at varying concentrations and for periods of time ranging from 1-96 hours showed that NPs were not associated with any effects on membrane integrity or membrane pore formation (Xu et al. 2021). However, there may be additional factors (food, mucus layer, etc) which modify *in vivo* responses from *in vitro* responses.

In addition to immune dysfunction, the ileum also showed alteration in other aspects of endocrine, cell metabolism and regulation. It is noteworthy that there were alterations in the expression of several genes implicated in peripheral thyroid hormone metabolism including deiodinase type 1 and 3. While other genes involved in thyroid hormone metabolism were slightly altered, changes in deiodinase have previously been reported as important targets to understand effects on thyroid hormone action (Amereh et al. 2020).

Reports of alterations in the expression of specific junctional or barrier proteins in the intestine are contradictory. There have been reports of altered junctional proteins following NP exposure in several species. These have frequently been decreased expression of proteins such as occludin, tight junction protein 1 (TJP1) also known as zona occludens, and some members of the claudin (CLDN) family of tight junction proteins. For example, Ma et al (Ma et al. 2024) reported decreased expression of CLDN3, CLDN5, and CLDN15 in mice injected with NPs. Liu et al (Liu et al. 2025) exposed mice for 10 weeks to both acrylamide and NPs via drinking water and observed decreased mRNA levels of CLDN5 and TJP1 in the colon. Li et al (Li et al. 2024) also exposed mice to PS NPs via drinking water for 6 weeks and reported unchanged protein levels of TJP1 and CLDN1 in the 0.1 mg/L group, but decreased protein levels of TJP1, CLDN1, and occludin in the 1, and 10 mg/L groups. Gosselink et al (Gosselink et al. 2026) exposed primary bronchial epithelial cell cultures to NPs and NPs and reported slightly decreased expression of TJP1 and occludin but did not observe any notable inflammation or cytotoxicity. A study similar to the current study was performed in which 100 nm NPs were administered via daily gavage to mice for 28 days, at a concentration of 1 mg/day (Xu et al. 2021). The doses of NPs ranged from 30-480 µg/ml. The authors reported effects on junction-related proteins in the intestinal epithelium which included a decrease in occludin and TJP1 but an upregulation of MMP-9. Comparable results on occludin and TJP1 were also reported in the mouse colon, in a study using NPs of 200 nm, 1 or 5 µm, administered by daily gavage to mice for 28 days (Zeng et al. 2024).

A decrease in CLDN1 staining was also observed. These results are comparable to those of Chen et al (Chen et al. 2024) who noted a decrease in CLDN1, occludin, and TJP1 expression (in intestinal epithelium) in mice given NPs by gavage for 21 days. Su et al (Su et al. 2024) also observed decreased expression of CLDN1 in the duodenum and jejunum of mice given MPs for 28 days via feeding (MPs incorporated into food pellets), but an increase in CLDN1 expression in the ileum. In the same experiments, these authors observed a similar pattern of expression with regards to several other tight junction proteins, including occludin, CLDN-2, −7, and −15 (jejunum only). The authors interpreted these results as suggesting that the increased expression of these claudins resulted in an increase in intestinal permeability. It should be noted that within the claudin family of proteins, some members are associated with forming a tight junctional barrier (CLDNs-1, −3, −5, −11, −18), while others form pores within the tight junctional complex and are thus associated with increased permeability(Gunzel and Yu 2013; Lee 2015). Similarly, an *in vitro* study by Jin and colleagues (Jin et al. 2019), in which Caco-2 cells were exposed to 5 µm MPs for 24 hrs, reported a down-regulation of CLDN1, as well as in occludin and TJP1, both associated with cellular junctions. Comparable observations were also reported by Liang et al (Liang et al. 2021) using a single oral dose of NPs of various sizes. Paradoxically, Zeng et al (Zeng et al. 2024) also reported an increase in CLDN1 in *in vitro* experiments with Caco-2 cells exposed to 5 µM (100 µg/ml) of PS-NPs for 24 hrs. The majority of *in vivo* studies using NPs in the literature report observations of altered tight junction protein expression that contrast our observations, in which few changes were observed in CLDN1 staining in mice administered 500 nm PS-NP particles for 60 days via gavage. Furthermore, our RNA-Seq data suggested only a minor but not significant upregulation of CLDN1. There were also few consistent changes noted in CDH1 immunostaining.

In addition, CLDN3 and CLDN7 expression was only slightly altered in the intestines of mice treated with either LD or HD NPs (500 nm) for 60 days. CLDN3 is known to be highly expressed in the gut and is associated with gut barrier integrity (Ahmad et al. 2023; Lu et al. 2013; Patel et al. 2012).Several studies have shown a decrease in CLDN3 expression concomitant with gut dysbiosis in various mouse models of colitis, as well as in human patients who have inflammatory bowel disease (IBD) (Ahmad et al. 2017; Lu et al. 2013; Patel et al. 2012). Furthermore, no significant alterations in CLDN3 or CLDN7 levels were observed in the jejunum, although RNA Seq data did show downregulation of CLDN7. Numerous reports of intestinal dysfunction and alterations in the murine intestine following NP exposure include indirect consequences of changes to microbiota as well as direct effects to the intestinal epithelium resulting in increased pro-inflammatory cytokine levels (such as IL-1b, TGFb1) and ROS production (Zeng et al. 2024). *In vitro* studies using Caco-2 cells cultured for up to 72 hrs showed a small but significant decrease in TEER in the cells exposed to highest dose of PS-NPs. These support our *in vivo* observations that 500 nm PS-NPs exert a small effect on the tight junctions of the small intestine.

The regulation of barrier protein expression, including claudins, is complex and multifactorial. It is possible that the presence of multiple claudin family members throughout the intestinal tract might have allowed for some compensatory mechanisms in the event that expression of a particular claudin was altered due of NP exposure. It is also possible that any discrepancies between our data and previous studies may reflect differences in NP particle composition, size, route, dose, and duration of NP administration.

Nonetheless, the DEGs that were up- or down-regulated by 2-fold or more between PS-NP-exposed mice and controls were largely associated with immune responses or immune-related processes, metabolism, and to a lesser degree, junctional proteins. Our data support previous observations that NPs represent a serious toxicological and physiological concern regarding the health and functioning of cells, animals, or humans exposed to NPs.

## CONCLUSIONS

Our results, as well as most of the literature pertaining to NP exposure, particularly PS-NPs, regardless of *in vivo* or *in vitro* experiments, indicate that NPs seem to affect the immune system, either directly or indirectly, and induce ROS production and inflammation. In the intestine, alterations to the microbiota are also an important aspect of NPs effects. A variety of cytokines, including TNF-α, IL1, and IL6β, have been reported to increase following NP exposure. Our data support these observations that sub-chronic exposure to 500 nm PS-NPs trigger components of inflammation and immune responses. TNF-α and IL1 are also known to down-regulate CLDN3, one of the tight junction proteins examined in this study and associated with gut barrier integrity. Clearly, a complex interplay among several regulatory pathways exists between the junctional proteins, immune and inflammatory responses to PS-NP exposure. Given that NPs occur in a wide variety of products, as well as in a vast array of sizes and chemical composition, and that the primary route of exposure is ingestion, it is evident that further research is warranted on elucidating mechanisms of action of NPs and on assessing their impact to humans and other species.

## ACKNOWLEDGEMENTS

The authors wish to thank Icten Meras (INRS) for her assistance with Caco-2 cell cultures.

## DATA AVAILABILITY

The raw RNA sequencing data have been submitted to the NCBI BioProject under accession number (Pending)

---

**Supplemental Figure 1.**
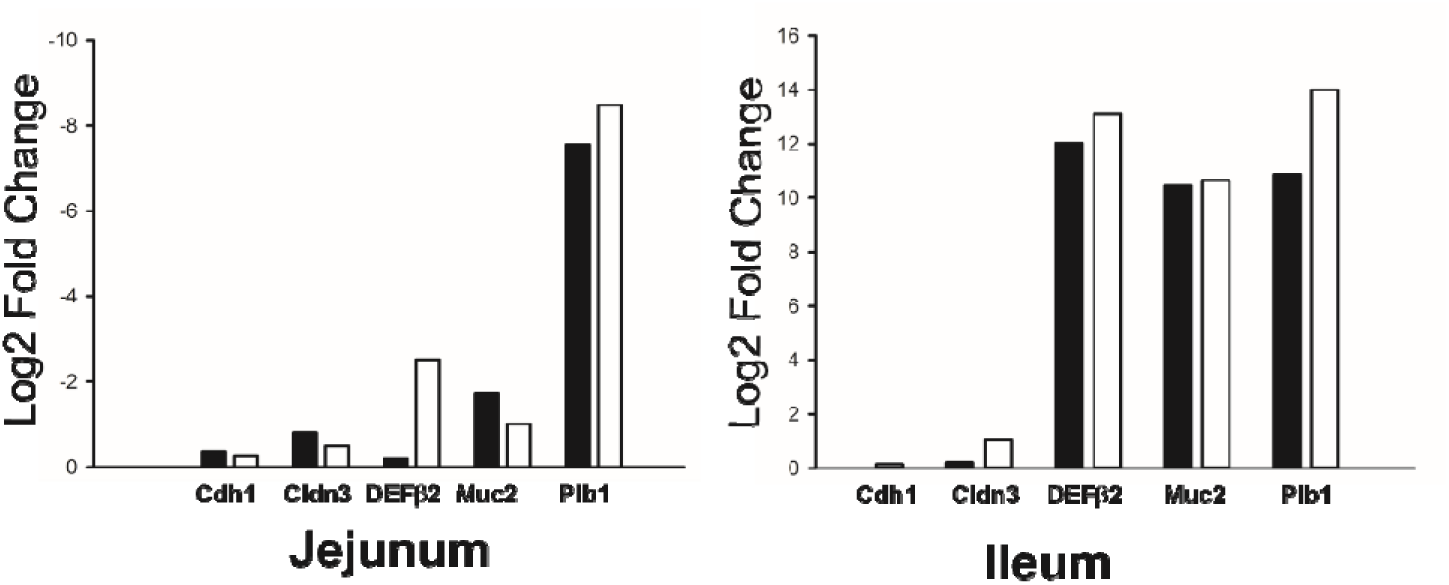
Select genes were verified by qPCR and log2 fold change for qPCR and for RNA seq were compared. Graphs represent values for qPCR (white bars) and for RNA Seq (black bars); note that genes for jejunum were downregulated while genes fur ileum were upregulated as compared to controls. Three (3) separate samples per group (control and PS-NP) were used for each analyses.

**Supplemental Table 1.** List of antibodies used in the study.

| Antibody Name | Manufacturer,<br>Catologue<br>Number | Concentration used in<br>immunohistochemistry<br>(µg/ml) | RRID |
| --- | --- | --- | --- |
| Cadherin 1 (E- | ABclonal |  |  |
| Cadherin) | A20798<br>Rabbit<br>polyclonal | 8.1 µg/ml | RRID:AB_3107194 |
| Musashi 1 | R&D/BioTechne<br>AF2628<br>Goat polyclonal | 2 µg/ml | AB_2147926 |
| Goat anti-rabbit IgG<br>HRP | Abcam ab6721 | 6 µg/ml | RRID:AB_955447 |
| Peroxidase-AffiniPure Donkey<br>anti-goat IgG<br>(H&L) | Jackson<br>Immunoresearch<br>705-035-003 | 1.6 µg/ml | AB_2340390 |

**Supplemental Table 2.**
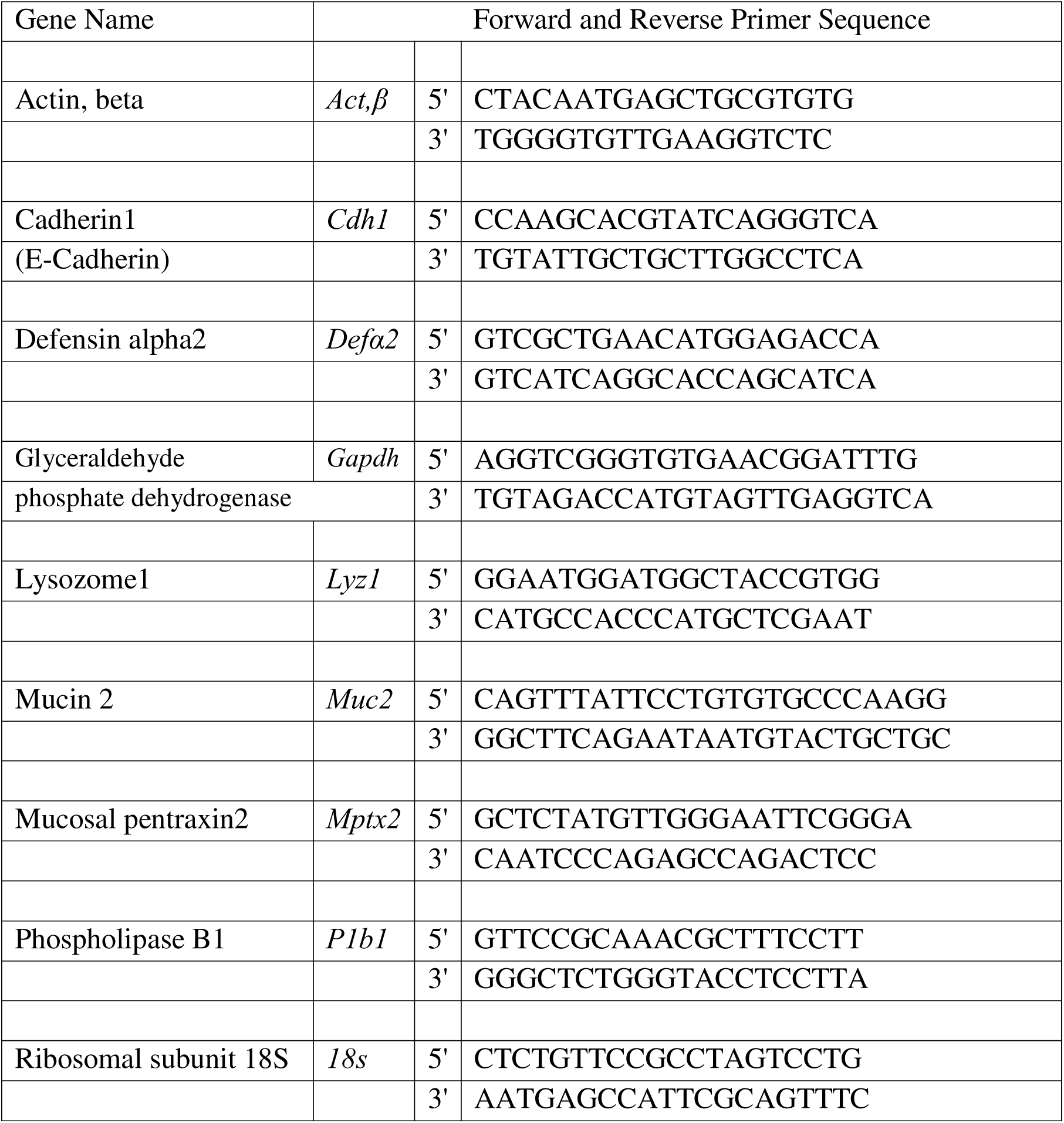
Sequence of qPCR primers used to validate RNAseq data.

**Supplemental Table 3.**
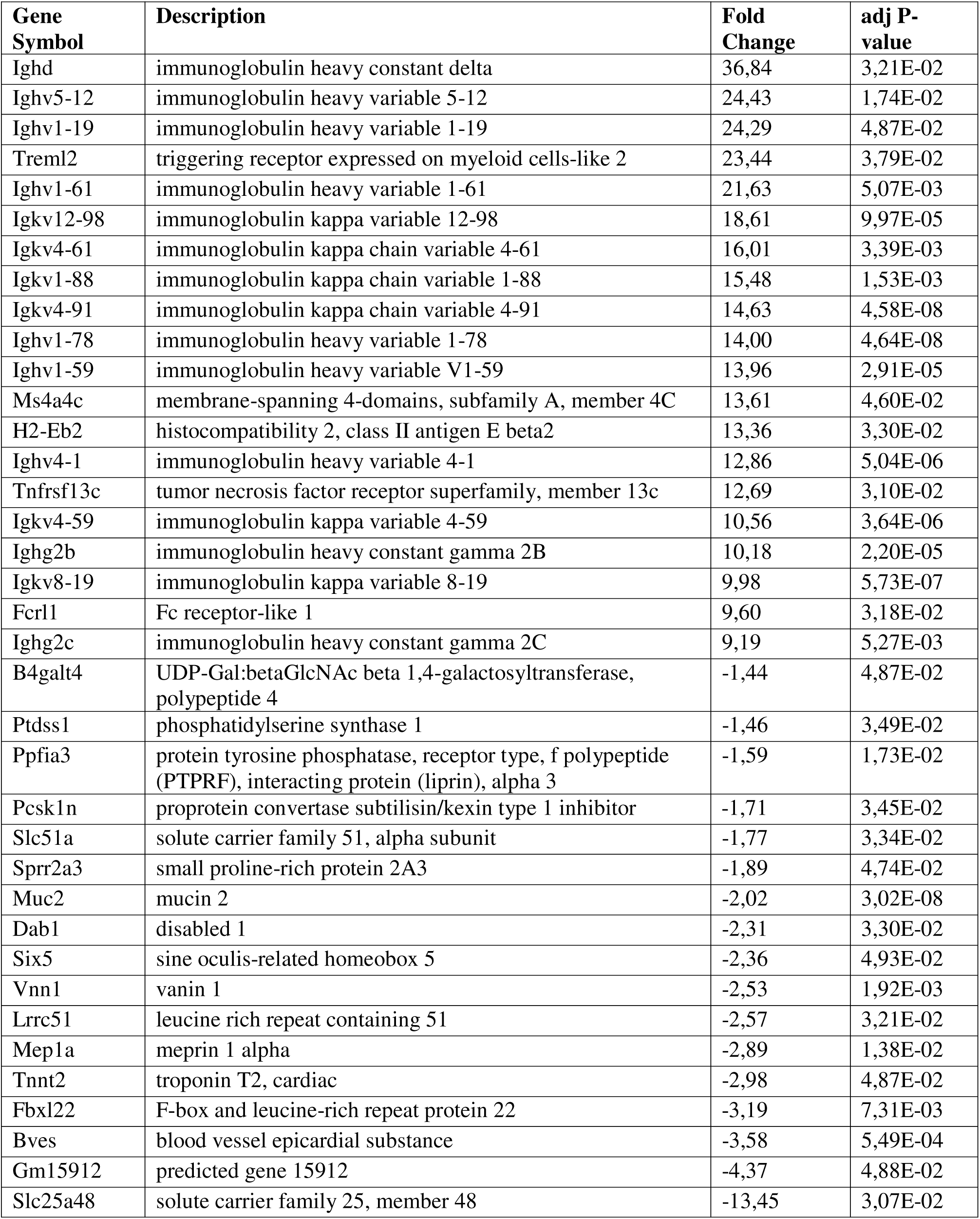
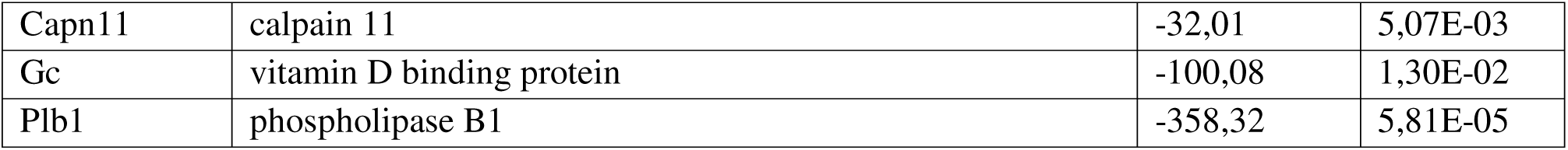
Top 20 up and downregulated DEGs in the jejunum of PS-NP treated mice versus control.

**Supplemental Table 4.**
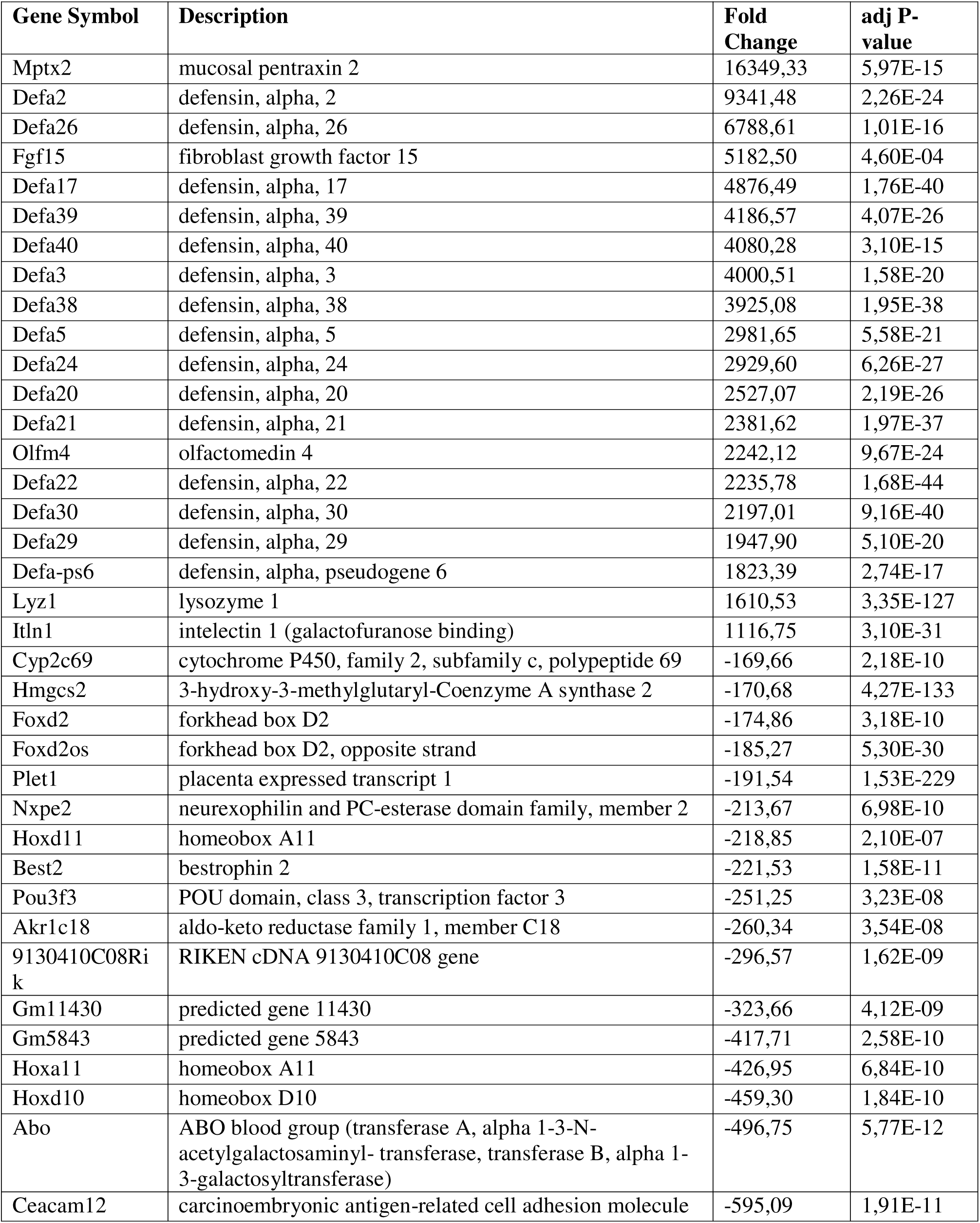
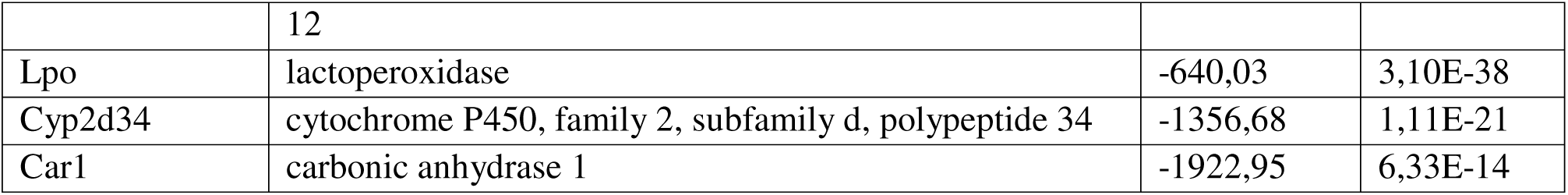
Top 20 up and down regulated DEGs in the ileum of PS-NP treated mice versus control.

| Gene Symbol | Description | Fold Change | adj P-value |
| --- | --- | --- | --- |
| Mptx2 | mucosal pentraxin 2 | 16349,33 | 5,97E-15 |
| Defa2 | defensin, alpha, 2 | 9341,48 | 2,26E-24 |
| Defa26 | defensin, alpha, 26 | 6788,61 | 1,01E-16 |
| Fgf15 | fibroblast growth factor 15 | 5182,50 | 4,60E-04 |
| Defa17 | defensin, alpha, 17 | 4876,49 | 1,76E-40 |
| Defa39 | defensin, alpha, 39 | 4186,57 | 4,07E-26 |
| Defa40 | defensin, alpha, 40 | 4080,28 | 3,10E-15 |
| Defa3 | defensin, alpha, 3 | 4000,51 | 1,58E-20 |
| Defa38 | defensin, alpha, 38 | 3925,08 | 1,95E-38 |
| Defa5 | defensin, alpha, 5 | 2981,65 | 5,58E-21 |
| Defa24 | defensin, alpha, 24 | 2929,60 | 6,26E-27 |
| Defa20 | defensin, alpha, 20 | 2527,07 | 2,19E-26 |
| Defa21 | defensin, alpha, 21 | 2381,62 | 1,97E-37 |
| Olfm4 | olfactomedin 4 | 2242,12 | 9,67E-24 |
| Defa22 | defensin, alpha, 22 | 2235,78 | 1,68E-44 |
| Defa30 | defensin, alpha, 30 | 2197,01 | 9,16E-40 |
| Defa29 | defensin, alpha, 29 | 1947,90 | 5,10E-20 |
| Defa-ps6 | defensin, alpha, pseudogene 6 | 1823,39 | 2,74E-17 |
| Lyz1 | lysozyme 1 | 1610,53 | 3,35E-127 |
| Itln1 | intelectin 1 (galactofuranose binding) | 1116,75 | 3,10E-31 |
| Cyp2c69 | cytochrome P450, family 2, subfamily c, polypeptide 69 | -169,66 | 2,18E-10 |
| Hmgcs2 | 3-hydroxy-3-methylglutaryl-Coenzyme A synthase 2 | -170,68 | 4,27E-133 |
| Foxd2 | forkhead box D2 | -174,86 | 3,18E-10 |
| Foxd2os | forkhead box D2, opposite strand | -185,27 | 5,30E-30 |
| Plet1 | placenta expressed transcript 1 | -191,54 | 1,53E-229 |
| Nxpe2 | neurexophilin and PC-esterase domain family, member 2 | -213,67 | 6,98E-10 |
| Hoxd11 | homeobox A11 | -218,85 | 2,10E-07 |
| Best2 | bestrophin 2 | -221,53 | 1,58E-11 |
| Pou3f3 | POU domain, class 3, transcription factor 3 | -251,25 | 3,23E-08 |
| Akr1c18 | aldo-keto reductase family 1, member C18 | -260,34 | 3,54E-08 |
| 9130410C08Rik | RIKEN cDNA 9130410C08 gene | -296,57 | 1,62E-09 |
| Gm11430 | predicted gene 11430 | -323,66 | 4,12E-09 |
| Gm5843 | predicted gene 5843 | -417,71 | 2,58E-10 |
| Hoxa11 | homeobox A11 | -426,95 | 6,84E-10 |
| Hoxd10 | homeobox D10 | -459,30 | 1,84E-10 |
| Abo | ABO blood group (transferase A, alpha 1-3-N-acetylgalactosaminyl- transferase, transferase B, alpha 1-3-galactosyltransferase) | -496,75 | 5,77E-12 |
| Ceacam12 | carcinoembryonic antigen-related cell adhesion molecule | -595,09 | 1,91E-11 |
|  | 12 |  |  |
| Lpo | lactoperoxidase | -640,03 | 3,10E-38 |
| Cyp2d34 | cytochrome P450, family 2, subfamily d, polypeptide 34 | -1356,68 | 1,11E-21 |
| Car1 | carbonic anhydrase 1 | -1922,95 | 6,33E-14 |

## REFERENCES

Ahmad R, Kumar B, Chen Z, Chen X, Muller D, Lele SM, Washington MK, Batra SK, Dhawan P, Singh AB. 2017. Loss of claudin-3 expression induces il6/gp130/stat3 signaling to promote colon cancer malignancy by hyperactivating wnt/beta-catenin signaling. Oncogene. 36(47):6592–6604.

Ahmad R, Kumar B, Thapa I, Talmon GA, Salomon J, Ramer-Tait AE, Bastola DK, Dhawan P, Singh AB. 2023. Loss of claudin-3 expression increases colitis risk by promoting gut dysbiosis. Gut Microbes. 15(2):2282789.

Amereh F, Babaei M, Eslami A, Fazelipour S, Rafiee M. 2020. The emerging risk of exposure to nano(micro)plastics on endocrine disturbance and reproductive toxicity: From a hypothetical scenario to a global public health challenge. Environ Pollut. 261:114158.

Andrady AL, Neal MA. 2009. Applications and societal benefits of plastics. Philos Trans R Soc Lond B Biol Sci. 364(1526):1977–1984.

Bashir ST, Chiu K, Zheng E, Martinez A, Chiu J, Raj K, Stasiak S, Lai NZE, Arcanjo RB, Flaws JA, Nowak RA. 2022. Subchronic exposure to environmentally relevant concentrations of di-(2-ethylhexyl) phthalate differentially affects the colon and ileum in adult female mice. Chemosphere. 309(Pt 1):136680.

Chakraborty S, Chakraborty A, Ghosh A, Bishwanath Singh N, Ghosh D, Kundu T, Das A. 2026. Microplastics and nanoplastics in human toxicity: Ros-mediated mechanisms, cellular damage and systemic effects. Mol Biol Rep. 53(1):1391.

Chen X, Xuan Y, Chen Y, Yang F, Zhu M, Xu J, Chen J. 2024. Polystyrene nanoplastics induce intestinal and hepatic inflammation through activation of nf-kappab/nlrp3 pathways and related gut-liver axis in mice. Sci Total Environ. 935:173458.

Choi H, Kaneko S, Suzuki Y, Inamura K, Nishikawa M, Sakai Y. 2024. Size-dependent internalization of microplastics and nanoplastics using in vitro model of the human intestine-contribution of each cell in the tri-culture models. Nanomaterials (Basel). 14(17):1435.

Cox KD, Covernton GA, Davies HL, Dower JF, Juanes F, Dudas SE. 2019. Human consumption of microplastics. Environ Sci Technol. 53(12):7068–7074.

Das A. 2023. The emerging role of microplastics in systemic toxicity: Involvement of reactive oxygen species (ros). Sci Total Environ. 895:165076.

Ding Y, Zhang R, Li B, Du Y, Li J, Tong X, Wu Y, Ji X, Zhang Y. 2021. Tissue distribution of polystyrene nanoplastics in mice and their entry, transport, and cytotoxicity to ges-1 cells. Environ Pollut. 280:116974.

Djouina M, Vignal C, Dehaut A, Caboche S, Hirt N, Waxin C, Himber C, Beury D, Hot D, Dubuquoy L et al. 2022. Oral exposure to polyethylene microplastics alters gut morphology, immune response, and microbiota composition in mice. Environ Res. 212(Pt B):113230.

Domenech J, de Britto M, Velazquez A, Pastor S, Hernandez A, Marcos R, Cortes C. 2021. Long-term effects of polystyrene nanoplastics in human intestinal caco-2 cells. Biomolecules. 11(10):1442.

Du B, Li T, He H, Xu X, Zhang C, Lu X, Wang Y, Cao J, Lu Y, Liu Y et al. 2024. Analysis of biodistribution and in vivo toxicity of varying sized polystyrene micro and nanoplastics in mice. Int J Nanomedicine. 19:7617–7630.

Eriksen M, Cowger W, Erdle LM, Coffin S, Villarrubia-Gomez P, Moore CJ, Carpenter EJ, Day RH, Thiel M, Wilcox C. 2023. A growing plastic smog, now estimated to be over 170 trillion plastic particles afloat in the world’s oceans-urgent solutions required. PLoS One. 18(3):e0281596.

Fan W, Xia D, Zhu Q, Hu L, Gan Y. 2016. Intracellular transport of nanocarriers across the intestinal epithelium. Drug Discov Today. 21(5):856–863.

Geyer R, Jambeck JR, Law KL. 2017. Production, use, and fate of all plastics ever made. Sci Adv. 3(7):e1700782.

Ghosh A, Bhakta S, Kapse N, Dhakephalkar PK, Patra C, Gorain B. 2026. Micro/nanoplastic-mediated gut dysbiosis and its impact on cardiac and neuroimmune function in zebrafish model: A multi-omics approach. Sci Total Environ. 1017:181443.

Gosselink IF, Leonhardt P, Drittij MJ, Hoppener EM, Smelt RJ, van Schooten FJ, Remels AHV. 2026. Evaluating the toxicity of polystyrene micro- and nanoplastics in human bronchial epithelial cells: Differences and challenges using aerosol and suspension exposures. Inhal Toxicol. 38(8):469–481.

Gunzel D, Yu AS. 2013. Claudins and the modulation of tight junction permeability. Physiol Rev. 93(2):525–569.

Hartmann NB, Huffer T, Thompson RC, Hassellov M, Verschoor A, Daugaard AE, Rist S, Karlsson T, Brennholt N, Cole M et al. 2019. Are we speaking the same language? Recommendations for a definition and categorization framework for plastic debris. Environ Sci Technol. 53(3):1039–1047.

He T, Qu Y, Yang X, Liu L, Xiong F, Wang D, Liu M, Sun R. 2023. Research progress on the cellular toxicity caused by microplastics and nanoplastics. J Appl Toxicol. 43(11):1576–1593.

Hirt N, Body-Malapel M. 2020. Immunotoxicity and intestinal effects of nano- and microplastics: A review of the literature. Part Fibre Toxicol. 17(1):57.

Hu CJ, Garcia MA, Nihart A, Liu R, Yin L, Adolphi N, Gallego DF, Kang H, Campen MJ, Yu X. 2024. Microplastic presence in dog and human testis and its potential association with sperm count and weights of testis and epididymis. Toxicol Sci. 200:235–240.

Huang J, Sun X, Wang Y, Su J, Li G, Wang X, Yang Y, Zhang Y, Li B, Zhang G et al. 2023. Biological interactions of polystyrene nanoplastics: Their cytotoxic and immunotoxic effects on the hepatic and enteric systems. Ecotoxicol Environ Saf. 264:115447.

Jin Y, Lu L, Tu W, Luo T, Fu Z. 2019. Impacts of polystyrene microplastic on the gut barrier, microbiota and metabolism of mice. Sci Total Environ. 649:308–317.

Kaur M, Sharma A, Bhatnagar P. 2024. Vertebrate response to microplastics, nanoplastics and co-exposed contaminants: Assessing accumulation, toxicity, behaviour, physiology, and molecular changes. Toxicol Lett. 396:48–69.

Kong L, Li S, Fu Y, Cai Q, Zhai Z, Liang J, Ma T. 2025. Microplastics/nanoplastics contribute to aging and age-related diseases: Mitochondrial dysfunction as a crucial role. Food Chem Toxicol. 199:115355.

Lee SH. 2015. Intestinal permeability regulation by tight junction: Implication on inflammatory bowel diseases. Intest Res. 13(1):11–18.

Li L, Lv X, He J, Zhang L, Li B, Zhang X, Liu S, Zhang Y. 2024. Chronic exposure to polystyrene nanoplastics induces intestinal mechanical and immune barrier dysfunction in mice. Ecotoxicol Environ Saf. 269:115749.

Liang B, Zhong Y, Huang Y, Lin X, Liu J, Lin L, Hu M, Jiang J, Dai M, Wang B et al. 2021. Underestimated health risks: Polystyrene micro- and nanoplastics jointly induce intestinal barrier dysfunction by ros-mediated epithelial cell apoptosis. Part Fibre Toxicol. 18(1):20.

Liu H, Li H, Yao X, Yan X, Peng R. 2025. Environmental nanoplastics induce mitochondrial dysfunction: A review of cellular mechanisms and associated diseases. Environ Pollut. 382:126695.

Liu S, Wu X, Gu W, Yu J, Wu B. 2020. Influence of the digestive process on intestinal toxicity of polystyrene microplastics as determined by in vitro caco-2 models. Chemosphere. 256:127204.

Livak KJ, Schmittgen TD. 2001. Analysis of relative gene expression data using real-time quantitative pcr and the 2(-delta delta c(t)) method. Methods. 25(4):402–408.

Lu L, Chen B. 2020. Biochar-amendment-reduced cotransport of graphene oxide nanoparticles and dimethyl phthalate in saturated porous media. Sci Total Environ. 705:135094.

Lu Z, Ding L, Lu Q, Chen YH. 2013. Claudins in intestines: Distribution and functional significance in health and diseases. Tissue Barriers. 1(3):e24978.

Ma S, Wang L, Li S, Zhao S, Li F, Li X. 2024. Transcriptome and proteome analyses reveal the mechanisms involved in polystyrene nanoplastics disrupt spermatogenesis in mice. Environ Pollut. 342:123086.

Magri D, Sanchez-Moreno P, Caputo G, Gatto F, Veronesi M, Bardi G, Catelani T, Guarnieri D, Athanassiou A, Pompa PP, Fragouli D. 2018. Laser ablation as a versatile tool to mimic polyethylene terephthalate nanoplastic pollutants: Characterization and toxicology assessment. ACS Nano. 12(8):7690–7700.

Mahler GJ, Esch MB, Tako E, Southard TL, Archer SD, Glahn RP, Shuler ML. 2012. Oral exposure to polystyrene nanoparticles affects iron absorption. Nat Nanotechnol. 7(4):264–271.

Mi H, Ebert D, Muruganujan A, Mills C, Albou LP, Mushayamaha T, Thomas PD. 2021. Panther version 16: A revised family classification, tree-based classification tool, enhancer regions and extensive api. Nucleic Acids Res. 49(D1):D394–D403.

Okano F, Harusato A, Nakanishi Y, Abo H, Etienne-Mesmin L, Kato M, Itoh Y. 2025. Oral exposure to micro- and nanoplastics generated from polyethylene terephthalate suppresses acute intestinal damage in vivo. J Hazard Mater. 498:139809.

Paget V, Dekali S, Kortulewski T, Grall R, Gamez C, Blazy K, Aguerre-Chariol O, Chevillard S, Braun A, Rat P, Lacroix G. 2015. Specific uptake and genotoxicity induced by polystyrene nanobeads with distinct surface chemistry on human lung epithelial cells and macrophages. PLoS One. 10(4):e0123297.

Patel RM, Myers LS, Kurundkar AR, Maheshwari A, Nusrat A, Lin PW. 2012. Probiotic bacteria induce maturation of intestinal claudin 3 expression and barrier function. Am J Pathol. 180(2):626–635.

Qiao R, Sheng C, Lu Y, Zhang Y, Ren H, Lemos B. 2019. Microplastics induce intestinal inflammation, oxidative stress, and disorders of metabolome and microbiome in zebrafish. Sci Total Environ. 662:246–253.

Schirinzi GF, Perez-Pomeda I, Sanchis J, Rossini C, Farre M, Barcelo D. 2017. Cytotoxic effects of commonly used nanomaterials and microplastics on cerebral and epithelial human cells. Environ Res. 159:579–587.

Su QL, Wu J, Tan SW, Guo XY, Zou DZ, Kang K. 2024. The impact of microplastics polystyrene on the microscopic structure of mouse intestine, tight junction genes and gut microbiota. PLoS One. 19(6):e0304686.

Sun J, Teng M, Zhu W, Zhao X, Zhao L, Li Y, Zhang Z, Liu Y, Bi S, Wu F. 2024a. Microrna and gut microbiota alter intergenerational effects of paternal exposure to polyethylene nanoplastics. ACS Nano. 18(27):18085–18100.

Sun R, Liu M, Xiong F, Xu K, Huang J, Liu J, Wang D, Pu Y. 2024b. Polystyrene micro- and nanoplastics induce gastric toxicity through ros mediated oxidative stress and p62/keap1/nrf2 pathway. Sci Total Environ. 912:169228.

Sutton SC, Hills RD, Jr. 2025. Role of nanoplastics in decreasing the intestinal microbiome ratio: A review of the scope of polystyrene. Toxics. 13(12):1036.

Szklarczyk D, Kirsch R, Koutrouli M, Nastou K, Mehryary F, Hachilif R, Gable AL, Fang T, Doncheva NT, Pyysalo S et al. 2023. The string database in 2023: Protein-protein association networks and functional enrichment analyses for any sequenced genome of interest. Nucleic Acids Res. 51(D1):D638–D646.

Tavakolpournegari A, Kannan U, Gregory M, Dufresne J, Costantino S, Lefrancois S, Cyr DG. 2026. Polystyrene nanoplastics induce transient microglial activation via endolysosomal retention in the mouse cortex. J Hazard Mater. 517:143485.

Walczak AP, Hendriksen PJ, Woutersen RA, van der Zande M, Undas AK, Helsdingen R, van den Berg HH, Rietjens IM, Bouwmeester H. 2015. Bioavailability and biodistribution of differently charged polystyrene nanoparticles upon oral exposure in rats. J Nanopart Res. 17(5):231.

Wu P, Huang J, Zheng Y, Yang Y, Zhang Y, He F, Chen H, Quan G, Yan J, Li T, Gao B. 2019. Environmental occurrences, fate, and impacts of microplastics. Ecotoxicol Environ Saf. 184:109612.

Xiao J, Jiang X, Zhou Y, Sumayyah G, Zhou L, Tu B, Qin Q, Qiu J, Qin X, Zou Z, Chen C. 2022. Results of a 30-day safety assessment in young mice orally exposed to polystyrene nanoparticles. Environ Pollut. 292(Pt B):118184.

Xie X, Deng T, Duan J, Xie J, Yuan J, Chen M. 2020. Exposure to polystyrene microplastics causes reproductive toxicity through oxidative stress and activation of the p38 mapk signaling pathway. Ecotoxicol Environ Saf. 190:110133.

Xu D, Ma Y, Han X, Chen Y. 2021. Systematic toxicity evaluation of polystyrene nanoplastics on mice and molecular mechanism investigation about their internalization into caco-2 cells. J Hazard Mater. 417:126092.

Yan L, Wang J, Cai X, Liou YC, Shen HM, Hao J, Huang C, Luo G, He W. 2024. Macrophage plasticity: Signaling pathways, tissue repair, and regeneration. MedComm (2020). 5(8):e658.

Yang Q, Dai H, Cheng Y, Wang B, Xu J, Zhang Y, Chen Y, Xu F, Ma Q, Lin F, Wang C. 2023. Oral feeding of nanoplastics affects brain function of mice by inducing macrophage il-1 signal in the intestine. Cell Rep. 42(4):112346.

Yin K, Wang D, Zhao H, Wang Y, Guo M, Liu Y, Li B, Xing M. 2021. Microplastics pollution and risk assessment in water bodies of two nature reserves in jilin province: Correlation analysis with the degree of human activity. Sci Total Environ. 799:149390.

Yong CQY, Valiyaveettil S, Tang BL. 2020. Toxicity of microplastics and nanoplastics in mammalian systems. Int J Environ Res Public Health. 17(5).

Yuvika, Sharma D, Sharma A. 2026. Molecular insights into physiological impact of micro- and nano-plastics on the digestive system and gut-brain axis. Comp Biochem Physiol C Toxicol Pharmacol. 304:110473.

Zeng G, Li J, Wang Y, Su J, Lu Z, Zhang F, Ding W. 2024. Polystyrene microplastic-induced oxidative stress triggers intestinal barrier dysfunction via the nf-kappab/nlrp3/il-1beta/mclk pathway. Environ Pollut. 345:123473.

Zhang Y, Hou B, Liu T, Wu Y, Wang Z. 2023. Probiotics improve polystyrene microplastics-induced male reproductive toxicity in mice by alleviating inflammatory response. Ecotoxicol Environ Saf. 263:115248.

Zhang Y, Jia Z, Gao X, Zhao J, Zhang H. 2024. Polystyrene nanoparticles induced mammalian intestine damage caused by blockage of bnip3/nix-mediated mitophagy and gut microbiota alteration. Sci Total Environ. 907:168064.

Zhang Zea. 2023. Continuous oral exposure to micro- and nanoplastics induced gut microbiota dysbiosis, intestinal barrier and immune dysfunction in adult mice. Environ Int. 182:108353.

Zhou Y, Zhou B, Pache L, Chang M, Khodabakhshi AH, Tanaseichuk O, Benner C, Chanda SK. 2019. Metascape provides a biologist-oriented resource for the analysis of systems-level datasets. Nat Commun. 10(1):1523.

Zhu A, Ibrahim JG, Love MI. 2019. Heavy-tailed prior distributions for sequence count data: Removing the noise and preserving large differences. Bioinformatics. 35(12):2084–2092.

